# An evolutionarily conserved stop codon enrichment at the 5’ ends of mammalian piRNAs

**DOI:** 10.1101/2021.10.27.464999

**Authors:** Susanne Bornelöv, Benjamin Czech, Gregory J Hannon

## Abstract

PIWI-interacting RNAs (piRNAs) are small RNAs required to recognize and silence transposable elements. The 5’ ends of mature piRNAs are defined through cleavage of long precursor transcripts, primarily by Zucchini (Zuc). Zuc-dependent cleavage typically occurs immediately upstream of a uridine. However, Zuc lacks sequence preference *in vitro*, pointing towards additional unknown specificity factors. We examined murine piRNAs and revealed a strong and specific enrichment of three sequences (UAA, UAG, UGA)— corresponding to stop codons—at piRNA 5’ ends. Stop codon sequences were also enriched immediately after piRNA processing intermediates, reflecting their Zuc-dependent tail-to-head arrangement. Further analyses revealed that a Zuc *in vivo* cleavage preference at four sequences (UAA, UAG, UGA, UAC) promotes 5’ end stop codons. This observation was conserved across mammals and possibly further. Our work provides new insights into Zuc-dependent cleavage and may point to a previously unrecognized connection between piRNA biogenesis and the translational machinery.

## INTRODUCTION

PIWI-interacting RNAs (piRNAs) are a class of small RNAs, 23-32 nucleotides (nt) in length. They are predominantly expressed in the gonads of most animals where they are loaded onto and guide PIWI-clade proteins to recognise complementary RNAs ^1–4^. Most piRNAs are transcribed from discrete genomic loci, termed piRNA clusters ^4,5^, are often complementary to transposable elements, and participate in the recognition and repression of such elements to safeguard genome integrity and fertility. Notable exceptions are the mouse pachytene piRNAs, which are derived from so-called ‘pachytene piRNA clusters’ that are expressed during spermatogenesis at the pachytene stage of meiosis ^1,2,6,7^. Pachytene piRNA clusters show no enrichment of transposon-complementary sequences but may regulate some protein-coding genes ^8–11^.

Processing of the piRNA cluster transcripts into piRNAs happens through two interconnected pathways: phased biogenesis and the ping-pong cycle ^5,12–15^. Ping-pong amplification relies on the slicer activity of the PIWI domain in cytoplasmic PIWI proteins, such as Aub and Ago3 in flies (*Drosophila melanogaster)* and MIWI and MILI in mouse. Once loaded with a piRNA guide, these proteins recognize and cleave complementary RNAs between position 10 and 11 of the guide ^5,13^. Ultimately, this piRNA-guided cleavage simultaneously degrades transposon transcripts and produces more piRNAs.

In contrast, phased piRNA biogenesis relies on the endonuclease activity of fly Zucchini (Zuc) or its mouse ortholog PLD6 to cleave piRNA precursor transcripts ^16–19^. An initial cleavage event through the ping-pong machinery or by a yet unknown trigger gives rise to an initial 5’ monophosphate end on the piRNA precursor ^14,15,20^ that is loaded onto a PIWI protein and transported to the outer mitochondrial membrane ^21,22^. There, Zuc cleaves the piRNA precursor downstream of the PIWI-protected footprint with the help of several co-factors ^21–25^, including the RNA helicase MOV10L1 (Armi in flies), whose ATPase activity is required for Zuc-mediated cleavage ^26^. This cleavage gives rise to two fragments: a pre-piRNA and a shortened precursor transcript with a new 5’ monophosphate. The remaining precursor is thought to yet again be loaded onto a PIWI protein and cleaved by Zuc. This repeated and step-wise processing gives rise to a set of phased pre-piRNAs, where the 3’ end of each pre-piRNA is immediately followed by the 5’ end of the next one ^14,15^. To give piRNAs their final length, murine pre-piRNAs are trimmed by the 3’-5’ exonuclease activity of PNLDC1 together with its co-factor TDRKH ^27–29^. Interestingly, no fly ortholog to PNLDC1 exists and Zuc-dependent piRNA length is increased only by an average of 0.5 nt in the absence the TDRKH ortholog Papi ^15,30^.

While much of the initial characterisation of piRNA biogenesis was done in flies and mouse, both ping-pong and phased biogenesis, as well as pre-piRNA trimming, has been observed across most animals ^31^. An unresolved question is why the Zuc-dependent phased cleavage happens with high selectivity immediately upstream of a uridine (U), giving the piRNAs their characteristic U at the 5’ end, the so-called 1U bias. Both fly Zuc and mouse PLD6 display endonuclease activity but no sequence specificity *in vitro* ^18,19^. The Piwi specificity loop was originally proposed to contribute to preferential binding of 1U-piRNAs, however, experimental manipulation of the Piwi specificity loop in flies had little to no impact on piRNA abundance and loading preferences and only revealed a weak repulsion of 1C piRNAs ^32^. The 1U bias must therefore be predominantly determined by the Zuc-dependent cleavage machinery, but the mechanism that creates this bias is yet to be identified ^32^.

Here we report a previously unrecognized enrichment of trinucleotide sequences corresponding to the three stop codons (UAA, UAG, UGA) at piRNA 5’ ends, with an enrichment around twice that of 1U alone. We show that this is a robust pattern across over a hundred mouse samples from a diverse set of developmental time points and conditions. Furthermore, this pattern is evolutionarily conserved across mammals and potentially more broadly. The stop codon enrichment is driven by piRNAs produced through Zuc-mediated cleavage. Our analyses releveled that Zuc-mediated cleavage preferentially occur at stop codons and UAC sequences, resulting in a specific enrichment of stop codon sequences in the final piRNA population. These sequences could represent a preference of a specificity co-factor for Zuc and correspond to stop codons purely by chance or alternatively could indicate that an uncharacterized mechanism connects the translational machinery to Zuc-mediated phased biogenesis. While additional work will be required to establish this connection and to determine its molecular basis, our work provokes a new hypothesis regarding how 5’ ends are determined during piRNA biogenesis.

## RESULTS

### Mouse piRNAs are enriched for stop codon sequences at their 5’ ends

While piRNA clusters are often challenging to analyse bioinformatically due to their high repeat content, mouse pachytene piRNA clusters are in large uniquely mappable and provide an excellent model to study piRNA biogenesis. A recent report focusing on 100 highly expressed pachytene piRNA clusters suggested that translating ribosomes are involved in defining piRNA 5’ ends ^33^. Based on this result, we hypothesised that stop codons should be depleted towards the 5’ ends of piRNAs to allow the translating ribosome to reach the piRNA 5’ end downstream region. However, upon reanalysing published data ^33^, we instead uncovered a strong and specific preference for the three stop codon sequences (UAA, UAG, UGA) at the 5’ ends of piRNAs mapping to pachytene piRNA clusters (Fig. 1a, left). While a 1U signature is expected for piRNAs produced through Zuc-dependent phased biogenesis, stop codon sequences were 2.14-fold enriched (95% confidence interval 2.04-2.23) compared to other U*nn* sequences (p=5e-6; two-sided one sample t-test). Moreover, enrichment of stop codon sequences was only observed directly at the 5’ end of piRNAs at position 1-3 (Fig. 1a), suggesting that it is not driven by a general over-representation of stop codons in pachytene piRNA clusters. Similar enrichment was observed across the whole length of piRNA precursor transcripts (Fig. S1a), suggesting that this is a general feature of pachytene piRNAs.

**Fig. 1.**
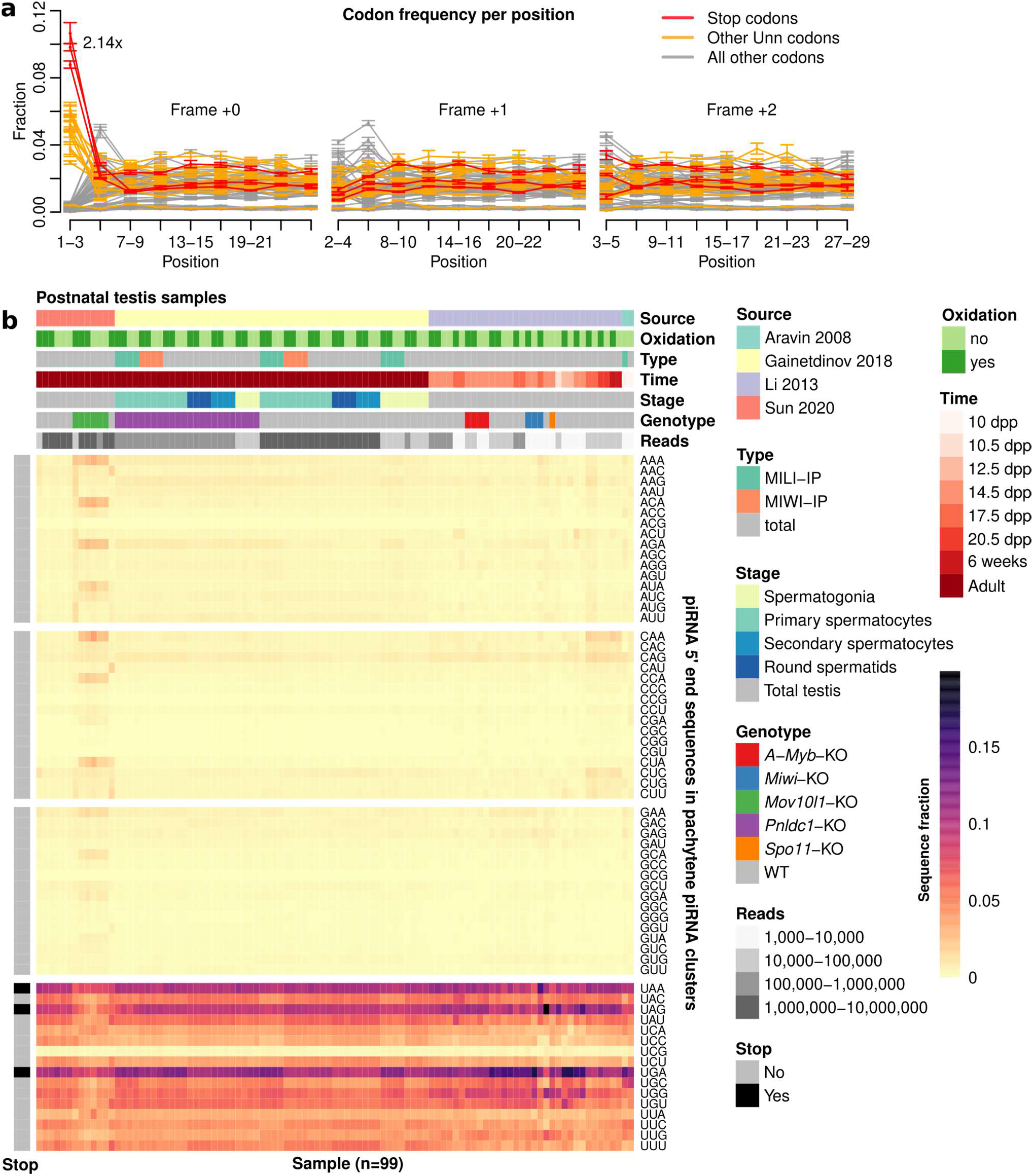
Postnatal pachytene piRNAs are enriched for stop codons at their 5’ ends. **a** Line graph showing the distribution of codons across piRNAs. Positions are numbered from the piRNA 5’ end. Frequency shown as mean ± one standard deviation (sd) (5 replicates). **b** Heatmap showing 5’ end sequence distribution of piRNAs mapping to pachytene piRNA clusters. Rows display relative sequence distribution and columns represent 99 sRNA-seq libraries derived from postnatal mouse testis. Column-wise annotations describe data source publication (Source), whether oxidated RNAs were captured (Oxidation), library type (Type), developmental time point (Time), spermatogenesis stage (Stage), mouse genotype (Genotype), and number of reads (Reads). Row-wise annotations show whether a sequence is a stop codon (Stop). Abbreviations: dpp, days post partum; KO, knockout; WT, wild-type. See also Fig. S1b for all piRNAs.

To exclude the possibility that this unexpected observation was the result of a technical artefact, we re-analysed 99 postnatal small RNA-seq libraries generated by different groups over a period of 12 years ^7,31,33,34^. Importantly, this included both oxidized and non-oxidized libraries from several knockout samples. Sequences corresponding to stop codons were enriched at piRNA 5’ ends across 61 investigated postnatal wild-type libraries (Fig. 1b). Moreover, despite a reduction in overall piRNA abundance, similar enrichment was observed in *A-Myb* (n=4), *Miwi* (n=3), *Spo11* (n=1), and *Pnldc1* knockout (n=24) libraries as well as for different immunoprecipitated libraries. These results suggest that the process favouring 5’ end stop codon sequences is independent of these factors, including MIWI binding to piRNA precursors. *Mov10l1* knockout libraries (n=6) reduced piRNA levels by 95 % ^33^ and displayed a slightly weaker enrichment of stop codons for the remaining piRNAs (Fig. 1b). This raises the possibility that Zuc-mediated cleavage—requiring the ATPase activity of MOV10L1 (ref ^26^)—promotes the enrichment of 5’ end stop codon sequences.

Pachytene piRNA clusters are non-coding transcripts and an enrichment of three specific codons (UAA, UAG, UGA) is therefore unexpected. Both adenosine (A) and guanosine (G) are purines and we therefore asked whether enrichment of stop codon sequences could be the result of an unrecognized U-purine-purine motif at piRNA 5’ ends. However, the fourth U-purine-purine sequence—UGG—is not enriched (Fig. 1a-b), and for simplicity we therefore refer to these three sequences as stop codons and to the over-representation of them compared with other U*nn* sequences at piRNA 5’ ends as a stop codon enrichment throughout this study, though we have not yet linked the function of these sequences as stop codons to piRNA biogenesis, per se.

We note that spermatogonia and some knockouts (*Mov10l1*, *A-Myb*, and *Spo11*) where spermatogenesis is arrested before the pachytene stage ^7^ are not expected to express many pachytene piRNAs. Moreover, a few libraries were excluded from our initial analysis as their piRNA count was below the cut-off of 1,000 reads. We therefore repeated the initial analysis without restricting it to pachytene piRNA clusters allowing 8 additional postnatal samples to be analysed (Fig. S1b), including *Mili* (n=1), *Dnmt1* (n=1) and *Pld6* (n=1) knockouts ^34,35^. Loss of PLD6 or MOV10L1 disrupted both 1U and stop codon enrichment on a global level (Fig. S1b), further indicating that 5’ end stop codons are a signature of the Zuc-mediated cleavage machinery. Similar observations were made when analysing piRNAs from postnatal ovaries (Fig. S2), where disruption of Zuc-mediated processing through *Mili* or *Pld6* knockout ^36^ or loss of oocytes through *Nanos3* knockout ^36^ resulted in a complete loss of both 1U and stop codon enrichment.

We next asked if stop codon enrichment could also be found in pre- and perinatal piRNAs. We used 25 small RNA-seq libraries ^34,35,37,38^ representing wild-type or *Tex15* knockout samples, which both harbour normal piRNAs ^37,38^, or *Pld6* knockout samples, which disrupt Zuc-mediated processing ^35^. Interestingly, a similar stop codon enrichment was observed also across pre- and perinatal samples, which was lost in the absence of PLD6 (Fig. 2a). Since pachytene piRNAs are produced though Zuc-mediated phased biogenesis, whereas pre- and perinatal piRNAs are also generated through ping-pong, we next restricted the analysis to piRNAs without an A at position 10, to exclude piRNAs with a clear ping-pong signature. This strengthened the stop codon enrichment, in particular in MIWI2-IP libraries (Fig. 2b), implicating Zuc-dependent phased biogenesis as a driver of stop codon enrichment also during embryonic development. Using published mappings and annotations ^37^, we next asked what classes of piRNAs show stop codon enrichment. Most piRNAs were repeat-derived (Fig. 2c) and piRNAs matching the sense strand of repeats had the strongest stop codon enrichment, followed by intergenic and intronic piRNAs (Fig. 2d).

**Fig. 2.**
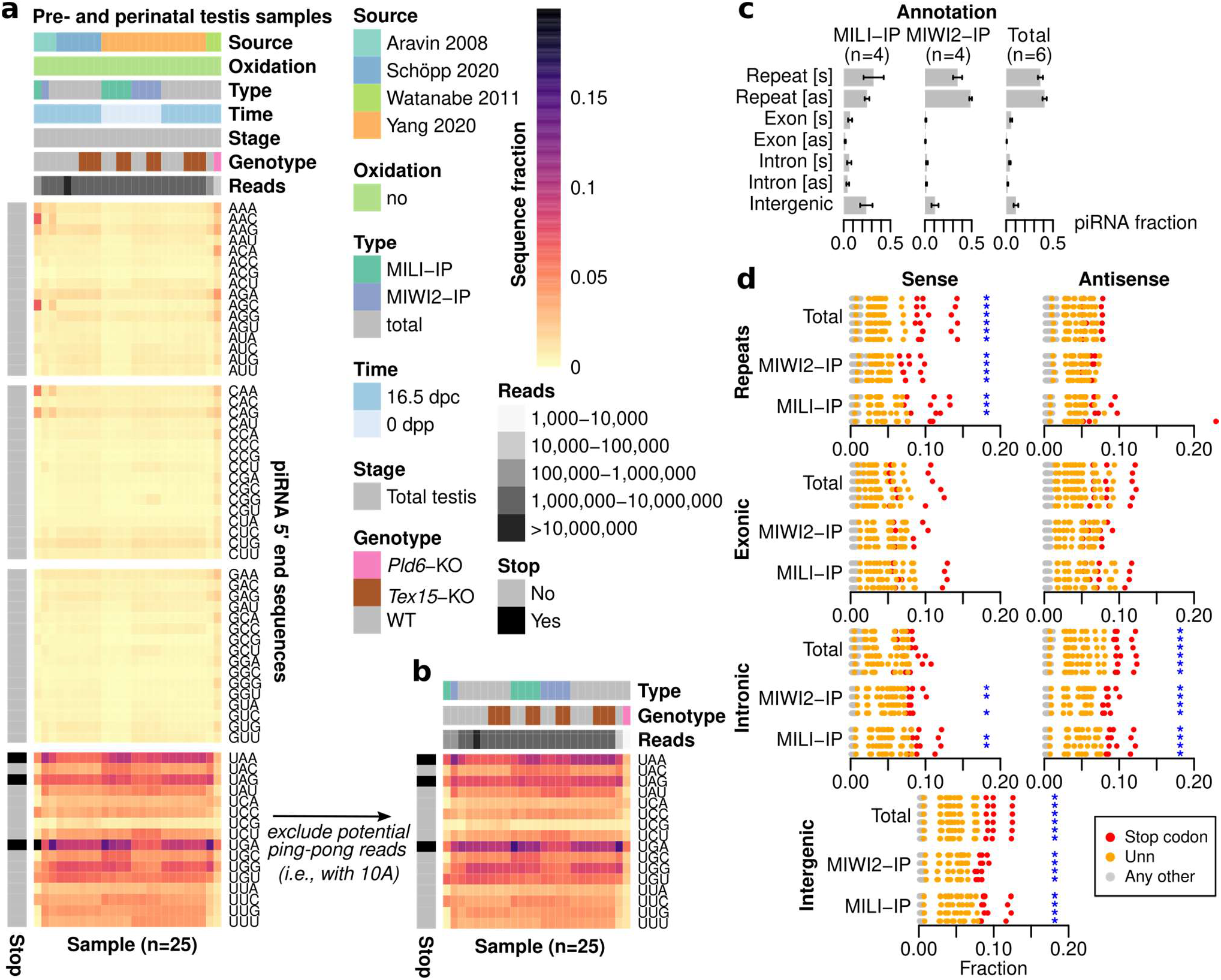
Pre- and perinatal piRNAs are enriched for stop codons at their 5’ ends. **a** Heatmap showing sequence distribution at piRNA 5’ ends across 25 small RNA-seq libraries from pre- and perinatal mouse testis. Each column represents one library. Column-wise annotations describe data source publication (Source), whether oxidated RNAs were captured (Oxidation), library type (Type), developmental time point (Time), spermatogenesis stage (Stage), mouse genotype (Genotype), and number of reads (Reads). Row-wise annotations show whether a sequence is a stop codon (Stop). Abbreviations: dpc, days post coitum; dpp, days post partum; WT, wild-type. **b** Heatmap showing re-processing of (a) using only reads without an A at position 10. **c** Annotation of piRNAs across 14 samples from Yang et al. ^37^ shown in (a). Bars represent mean fraction ±sd (4-6 replicates). Abbreviations: s, sense; as, antisense. **d** Overview of 5’ end sequences across the samples in (c) per annotation type. Each row represents one library (either WT or *Tex15*-KO). All possible 5’ end sequences are shown as circles and colour-coded to identify stop codons (*n*=3, red), other U*nn* sequences (*n*=13, orange), and all other sequences (*n*=48, grey). Libraries where the three stop codons are more abundant than any other sequence are marked with a blue asterisk.

This rank order may in part reflect signal robustness determined by piRNA abundance. No stop codon enrichment was observed for piRNAs antisense to repeats, consistent with their production predominantly through ping-pong ^34^. Moreover, we observed no clear differences in size between piRNAs with a 5’ stop codon and other piRNAs (Fig. S3), suggesting that 5’ end stop codons is a signature of bona fide piRNAs. Thus, searching for a translating ribosome signature across pachytene piRNAs, we instead detected a highly reproducible and unexpected enrichment of stop codons at piRNA 5’ ends. This enrichment appears to be unrelated to many piRNA factors and can be observed in both testis and ovary and across different developmental stages.

### Mouse pachytene pre-piRNAs show downstream stop codon enrichment

Mouse pachytene piRNAs are produced through Zuc-mediated phased biogenesis in which PLD6 (Zuc in flies) repeatedly cleaves the piRNA precursor. These cleavages give rise to phased pre-piRNAs, where each pre-piRNA 3’ end is immediately followed by the 5’ end of the next pre-piRNA ^14,15^. Pre-piRNAs are up to 50 nt long, with most of them being in the 29-33 nt range, and are trimmed to their mature length (26-27 nt) by PNLDC1 ^28,29^. Although pre-piRNAs occur only at very low levels in wild-type mice, PNLDC1-deficient mice show defective trimming, characterized by an accumulation of pre-piRNAs and a depletion of mature piRNAs ^28,29^. Trimmer mutants are characterized by a 1U signal downstream of pre-piRNA 3’ ends ^14,15^, and we hypothesized that this should be accompanied by a downstream stop codon enrichment if Zuc-mediated cleavage promotes both signals. We used small RNA-seq data from *Pnldc1* knockout mice ^31^ to test whether a stop codon enrichment was present both at piRNA 5’ ends and immediately downstream of pre-piRNA 3’ ends, consistent with the expected tail-to-head arrangement of pre-piRNAs produced by Zuc-dependent phased biogenesis.

As previously reported ^31^, libraries derived from *Pnldc1* knockout mice displayed an altered size profile (mostly 28-36 nt) consistent with defective trimming, whereas libraries from wild-type controls had a normal size distribution (mostly 26-31 nt) (Fig. 3a) with two peaks corresponding to piRNAs loaded onto MILI (26-27 nt) and MIWI (29-31 nt) (Fig. 3a, Fig. S4a). The *Pnldc1* knockout libraries also displayed a strong 1U bias as well as a stop codon enrichment immediately downstream of their 3’ ends, in sharp contrast to the wild-type libraries that displayed neither 1U nor any preference for stop codons after their 3’ ends (Fig. 3b). This strongly implies that the 5’ end stop codon enrichment is coupled to piRNA precursor cleavage and not to differences in (pre-)piRNA loading or stability. Notably, this analysis also excludes library preparation artefacts such as ligation biases ^39^ as a source for the observed stop codon enrichment.

**Fig. 3.**
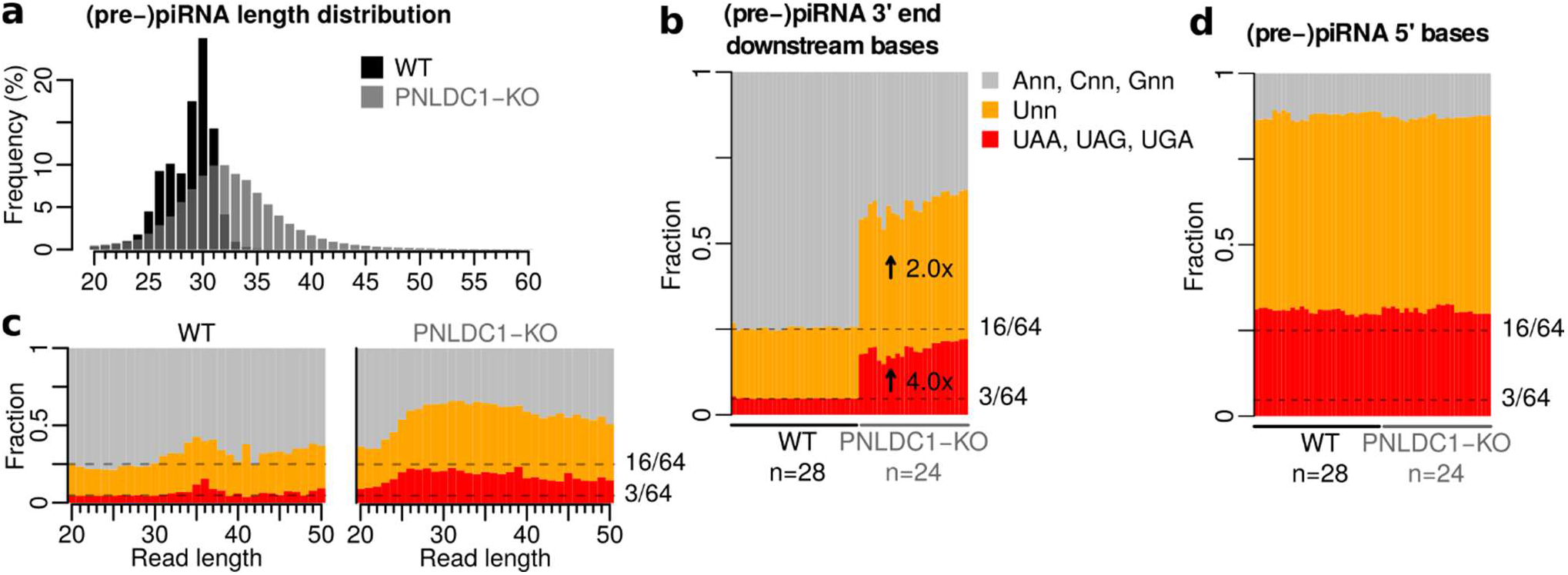
Stop codons are enriched immediately downstream of pachytene pre-piRNA 3’ ends. **a** Length distribution of PNLDC1 knockout (KO) and wild-type (WT) controls. Individual conditions shown in Fig. S4. **b** 1U and stop codon fraction immediately downstream of 3’ ends in wild-type (WT) and PNLDC1 knockout (KO) sRNA-seq libraries. Arrows indicate mean enrichment in KO relative to WT. Dashed lines indicate expected fraction assuming all trinucleotide sequences are equally abundant. **c** Bar graph showing mean fraction in (b) separated per read length across all WT (left) or PNLDC1-KO (right) sRNA-seq libraries. Individual conditions shown in Fig. S4. **d** 1U and stop codon fraction at the (pre-)piRNA 5’ ends in wild-type (WT) and PNLDC1-KO sRNA-seq libraries. Dashed lines indicate expected fraction assuming all trinucleotide sequences are equally abundant.

Further separating (pre-)piRNAs by their length revealed that while most (pre-)piRNAs in wild-type samples—including the by far most abundant lengths 26-27 nt—did not show any 1U or stop codon enrichment downstream of their 3’ ends, there was a rare population of reads with lengths of 35-36 nt with both a 1U and a stop codon enrichment downstream of the mature piRNAs (Fig. 3c, left). Considering their length and 1U bias, we conclude that these likely represent pre-piRNAs captured at low frequency under wild-type conditions. In contrast, *Pnldc1* knockout mice displayed strong 1U and stop codon enrichment for all read lengths, with only a minor reduction in the signal at the very short 20-22 nt lengths (Fig. 3c, right).

As an additional control, we compared the 1U and stop codon enrichment at the piRNA 5’ ends, revealing no difference between *Pnldc1* knockout and control mice (Fig. 3d). We note that both the 1U and stop codon enrichment was stronger at piRNA 5’ ends compared to downstream of pre-piRNA 3’ ends (Fig. 3b, d). This may be due to additional biases introduced through loading of the (pre-)piRNAs onto a PIWI protein ^32^ or due to incomplete processing of the precursor transcripts.

Thus, our results support a model in which mouse pachytene piRNAs are characterized by stop codons at the 5’ end and argue against alternative technical explanations for this observation. Furthermore, the 5’ end stop codon enrichment is driven by Zuc-mediated phased biogenesis and appears to co-occur with the 1U bias.

### A distinct cleavage preference promotes 5’ stop codons

To investigate to what extent nucleotide sequence contributes to 5’ end definition of mouse pachytene piRNAs, we developed a 5’ end definition score (Fig. 4a, see Methods for details). This score describes how likely each position within pachytene piRNA clusters is to be selected as a piRNA 5’ end compared with its neighbouring positions. Importantly, this score therefore reflects cleavage site selection directly and makes it independent of the underlying abundance of each sequence within the clusters. We reasoned that if there is an evolutionary advantage to piRNA 5’ end formation at stop codon sequences, we should detect a sequence specificity that not only promotes cleavage at stop codons, but also disfavours cleavage at most alternative sequences. This specificity could be expected to act strongly against abundant alternative sequences, whereas rare alternative sequences might be tolerated as long as they do not distort the global cleavage at stop codons.

**Fig. 4.**
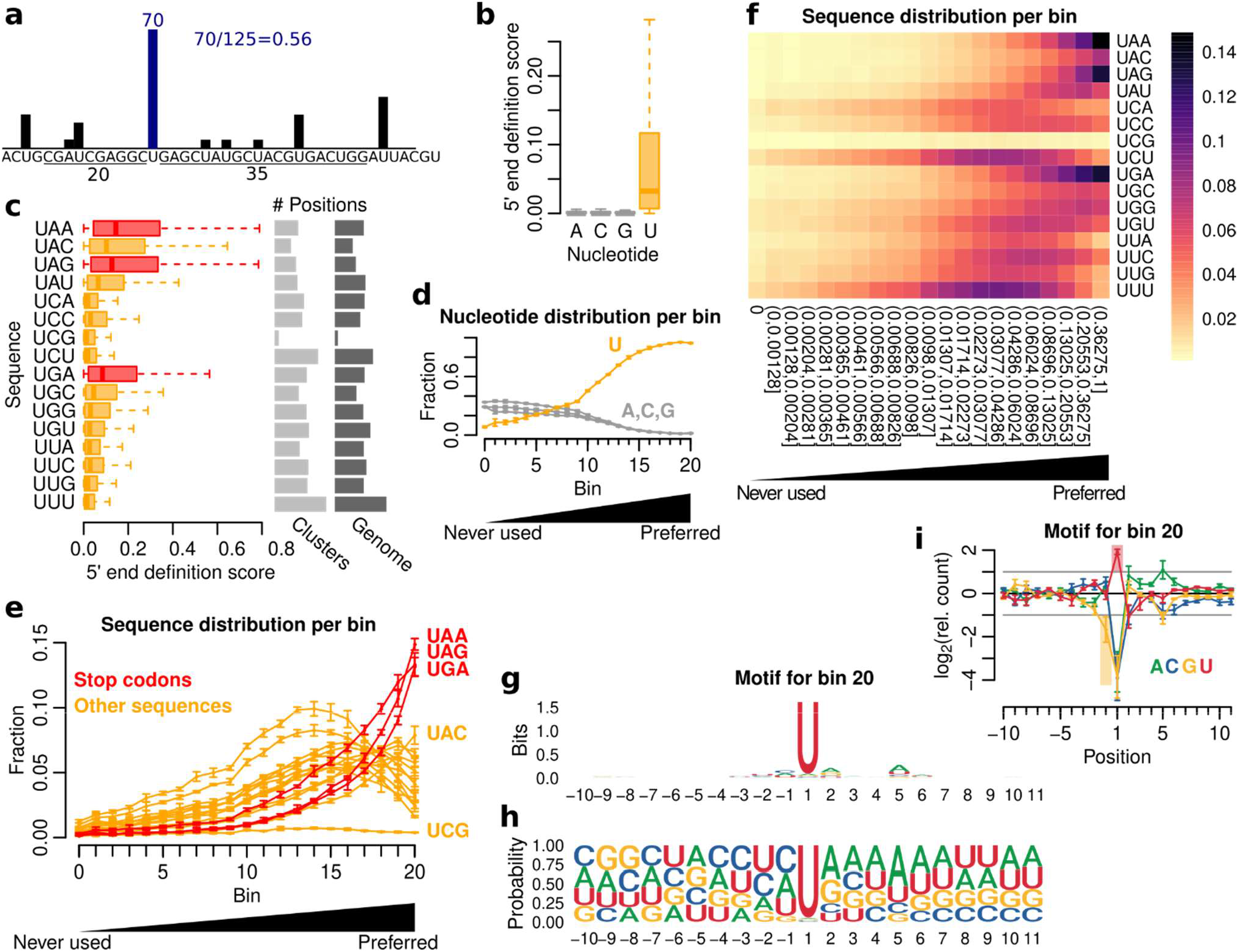
Deciphering the cleavage preference of PLD6. **a** Illustration of the 5’ end definition score. **b** Boxplot showing 5’ end definition score per nucleotide using pooled data (5 replicates). **c** Boxplot showing 5’ end definition score per U*nn* sequence. Stop codons are shown in red and the remaining sequences in orange. The number of positions per sequence is shown to the right. The data was pooled (5 replicates). **d** Line graph showing nucleotide fraction per bin. Fraction shown as mean ±sd (5 replicates). **e** Line graph showing U*nn* sequence fraction per bin. Fraction shown as mean ±sd (5 replicates). **f** Heatmap showing 5’ end sequence distribution per bin. The 5’ end definition score thresholds are shown under each column. Fraction calculated as mean across 5 replicates. An extended figure with all sequences is available as Fig. S5a. **g** Sequence motif around positions from bin 20 (pooled signal, 5 replicates). **h** Nucleotide frequency around positions from bin 20 (pooled signal, 5 replicates). **i** Nucleotide enrichment around positions from bin 20 (pooled signal ±sd, 5 replicates). An extended figure with all bins is available as Fig. S5b. Boxplots show median (central line), interquartile range (IQR, box), and minimum and maximum values (whiskers, at most 1.5*IQR). Additional analyses using a second dataset are available in Fig. S6 and Fig. S7.

The resulting scores were distributed between 0 (never observed as a 5’ end) and 1 (always selected as a 5’ end) with a mean of 0.0329. As expected, positions with a U nucleotide had considerably higher scores than other nucleotides (Fig. 4b), consistent with the known cleavage preference upstream of U. Notably, positions with a U as the first base of a stop codon scored higher than positions with a U in other trinucleotide contexts (Fig. 4c, left), indicating that stop codon enrichment at piRNA 5’ ends is driven by cleavage site preferences rather than an overrepresentation of stop codons in piRNA clusters. One minor exception from this pattern is the UAC sequence, which scored comparably to the lowest scoring stop codon (UGA). We speculate that UAC, which is similar in sequence to two of the stop codons, may also be tolerated by the cleavage machinery. From an evolutionary perspective, this may have been facilitated by its rare abundance as the second least frequent trinucleotide sequence (Fig. 4c, right). We therefore concluded that a distinct cleavage preference at four sequences promotes stop codon enrichment.

To study the dynamics in the selection of cleavage position for different sequence contexts, we next divided all cluster positions into 21 bins. The first bin represented 995,160 positions where no piRNA 5’ end was observed in any library, and the remaining 904,909 positions were divided into 20 equally sized bins ordered from lowest to highest scores. Thus, our 21 bins reflect the least to the most favourable cleavage positions, while correcting for the neighbouring sequence context and local piRNA abundance.

Notably, while bin 0 was strongly depleted for U at the 5’ ends, the following bins displayed a gradual increase in 1U enrichment, up to the last three bins, which were almost exclusively populated by positions with a 1U (Fig. 4d). Moreover, the sequence context for these Us revealed a striking enrichment of the three stop codon sequences specifically at the most preferred bin 20 (Fig. 4e-f). In contrast, other U*nn* sequences displayed their strongest enrichment earlier, at bin 14-19 (Fig. 4e-f), suggesting that while sequences starting with a U are preferred over non-U ones, stop codon sequences are specifically associated with the most preferred piRNA 5’ ends.

Sequences with multiple Us were generally enriched in lower bins compared with codons with a single U, likely reflecting competition between neighbouring Us in the selection of a cleavage site. However, this does not explain why stop codon sequences are favoured, since there are six other trinucleotide sequences with only one U and five of them have their highest enrichment earlier in bin 17-19 (Fig. 4f), while the sixth one (UAC) occurs only at low frequency in pachytene piRNA clusters.

We also noted that all sequences with CpG dinucleotide (including UCG) were strongly depleted (Fig. 4e, Fig. S5a), an observation that is in line with the genome-wide depletion of CpG dinucleotides in non-coding regions due to their high mutation rate associated with DNA methylation ^40^.

Notably, BmZuc—the silkmoth (*Bombyx mori*) ortholog to Zuc—was recently suggested to cleave piRNA precursors at a specific sequence motif ^41^. To perform a similar analysis in mouse, we derived a sequence motif for bin 20, representing the most preferred piRNA 5’ end positions (Fig. 4g-i). Aside from the expected 1U signal, the most striking pattern was that G nucleotides were strongly depleted immediately upstream of the preferred cleavage sites (Fig. 4i), as previously observed in both mouse and silkmoth ^41^. Interestingly, we also observed a GC-rich region further upstream of the cleavage position (Fig. 4h, i), in contrast to the AU-rich region downstream of the cleavage site (Fig. 4h, i). Moreover, nucleotide enrichments calculated for the other bins revealed no GC-rich upstream region (Fig. S5b), suggesting that a GC-rich upstream region may contribute to the definition of the most preferred cleavage sites.

To confirm that the preference for UAA, UAG, UGA and UAC sequences was driven by cleavage site selection during phased biogenesis, we repeated the above analysis using 52 additional libraries representing either wild-type or *Pnldc1* knockout testis ^31^. We reasoned that if Zuc-mediated phased cleavage gives rise to the observed cleavage site selection, then positions immediately downstream of pre-piRNAs 3’ ends in *Pnldc1* knockout— representing the next 5’ end—should display a similar signal. Cluster positions were separately scored against their background based on either (pre-)piRNA 5’ or 3’ ends as illustrated in Fig. S6a. As previously, Us were preferentially selected as 5’ ends positions (Fig. S6b), but only Us downstream of pre-piRNAs 3’ ends (in *Pnldc1* knockout) displayed higher scores compared to other nucleotides (Fig. S6b, right). Moreover, Us in stop codon or UAC context were preferred over Us in other U*nn* contexts both at 5’ ends and downstream of pre-piRNA 3’ ends (Fig. S6c), and pre-piRNA 5’ and 3’ end scores were highly correlated (Fig. S6d, r=0.95, p=1e-33). Finally, the most frequently used positions in bin 20 using either 5’ ends or pre-piRNA 3’ ends corresponded to stop codons (Fig. S7). Altogether, this confirms that stop codon enrichment is primarily driven by a cleavage preference at four separate sequences.

### Open reading frames do not contribute to piRNA 5’ end definition

The above analysis has shown that piRNA 5’ ends are preferably located at stop codon sequences. We reasoned that if piRNA production is directly triggered by the translational machinery, then stop codons that are part of an open reading frame (ORF) may have a different 5’ end definition score compared with those that are not. To test this idea, we identified 92,481 stop codons present in mouse pachytene piRNA precursor transcripts. Out of these, 22,374 are part of an ORF of length 3-100 amino acids (i.e., having an in-frame upstream start codon before any inframe upstream stop codon) and 59,227 are not. However, we observed no difference in piRNA 5’ end definition score between the two groups (Fig. S8a, top; p=0.50, Wilcoxon rank sum test). Similarly, no differences were observed when limiting the analysis to ORFs of certain size ranges (Fig. S8a, bottom). Therefore, the presence or absence of an ORF does not globally contribute to defining piRNA 5’ ends at stop codon positions.

Since the vast majority of ORFs are never translated, we next focused on ORFs with experimental evidence of translation from either ribosome profiling or proteomics ^42^. As shown in Fig. S8b, we observed no difference in the mean 5’ end definition score between stop codons from experimentally confirmed ORFs (0.135) and control stop codons (0.194; 95% CI 0.105-0.299; p=0.12; one-sided empirical test). Taken together, we found no evidence that the stop codon enrichment is connected to translated ORFs within the long non-coding transcripts.

### Cleavage preference is weakly mirrored by differences in sequence conservation

We next asked whether genomic positions that preferentially give rise to piRNA 5’ ends display higher conservation compared with non-preferred positions. We reasoned that if such a pattern of purifying selection is detected, this indicates that piRNAs with stop codon at their 5’ ends also have an important downstream function, whereas its absence instead supports the idea that 5’ end stop codons are simply an incidental signature of uncharacterized factors involved in Zuc-mediated cleavage. An initial analysis across our previously defined bins revealed no conservation signal at bin 0, a weak but gradually increasing conservation across bin 1-16, followed by a sharp reduction in conservation at bin 17-20 (Fig. 5a).

**Fig. 5.**
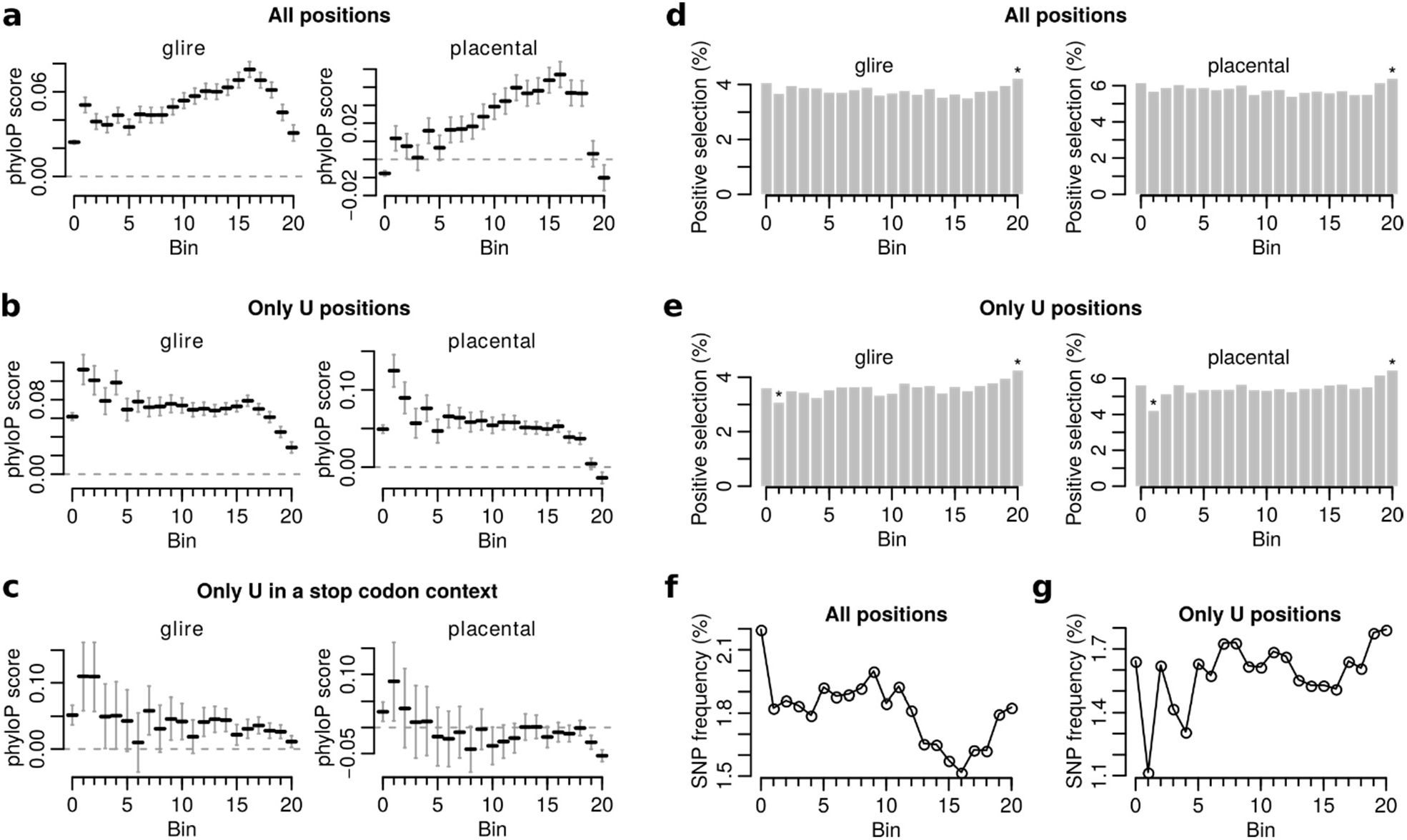
Sequence conservation at mouse pachytene piRNA clusters. **a** phyloP conservation scores (in glires and placental mammals, respectively) at cluster positions binned by piRNA 5’ end definition score. Horizontal lines represent the mean with a 95% confidence interval. **b** Same as (a), but only positions with a U nucleotide are shown. **c** Same as (a), but only positions with a U nucleotide in stop codon context are shown. **d** Bar plots showing the percentage of positions per bin with a significant phyloP score (in glires and placental mammals, respectively) indicating positive selection. Asterisks mark bins that were significantly different from all bins (p<0.05; two-sided Z test). **e** Same as (d), but only positions with a U nucleotide are shown. **f** SNP frequency at cluster positions binned by piRNA 5’ end definition score. A SNP was considered to be present at a certain position if it was listed in dbSNP. **g** Same as (f), but only positions with a U nucleotide are shown.

Except for the reduced conservation at the most preferred positions (Fig. 5a), this largely reflected the fraction of U positions in each bin (Fig. 4d). Since genomic A and T positions (A and U in the transcribed sequence) show higher conservation than C and G positions, we next restricted the analysis to U positions within each bin. The modified analysis revealed a steady level of weak conservation across all bins, except for bin 19-20 that displayed reduced levels of conservation (Fig. 5b). Further restricting the analysis to positions with a U in a stop codon context gave largely similar results (Fig. 5c). Notably, the drop in conservation score at bin 20 was mirrored by a higher number of positions showing signs of positive selection (Fig. 5d-e).

These observations were further supported by mouse variation data from dbSNP, revealing a similar pattern with decreasing SNP frequency across bin 1-16, followed by increased frequency at bin 17-20 (Fig. 5f). As previously, restricting the analysis to U positions resulted in a steady SNP frequency across most bins with the highest frequencies observed at bin 19-20 (Fig. 5g).

In summary, we did not find signs of purifying selection acting on Us in a stop codon context, suggesting that despite their enrichment at piRNA 5’ ends, stop codon sequences may not have a downstream function. Instead, the most preferred cleavage positions (bin 20) exhibited slightly less conservation and higher variability within murine piRNAs, likely reflecting a higher fraction of fast-evolving sequences such as transposons.

### Stop codon enrichment is conserved across mammals

Next, we asked whether the stop codon enrichment at the piRNA 5’ ends is specific to murine sequences. To this end, we used the piRNA cluster database ^43^ that includes 218 small RNA libraries from testes or ovaries across 49 species. In total, 33 species were represented with at least one testis library and 34 with at least one ovary library (see Supplementary Table S1). Consistent with their piRNA composition, nearly all libraries showed a strong preference for 1U (Fig. 6a), which increased further when we filtered the libraries to only include reads mapping to piRNA clusters (Fig. 6b). For each of the 211 libraries kept after filtering, we calculated the frequency of each trinucleotide sequence across its piRNA 5’ ends. We next calculated stop codon ratio as the ratio between stop codons and other U*nn* sequences at piRNA 5’ ends (see Methods). Notably, 176 out of 211 libraries (83%) showed an overrepresentation of stop codons over other U*nn* sequences. (Fig. 6c). This was particularly striking among mammals (74 out of 80 libraries; 93%) and ray-finned fish (36 out of 37 libraries; 97%) (Fig. 6c).

**Fig. 6.**
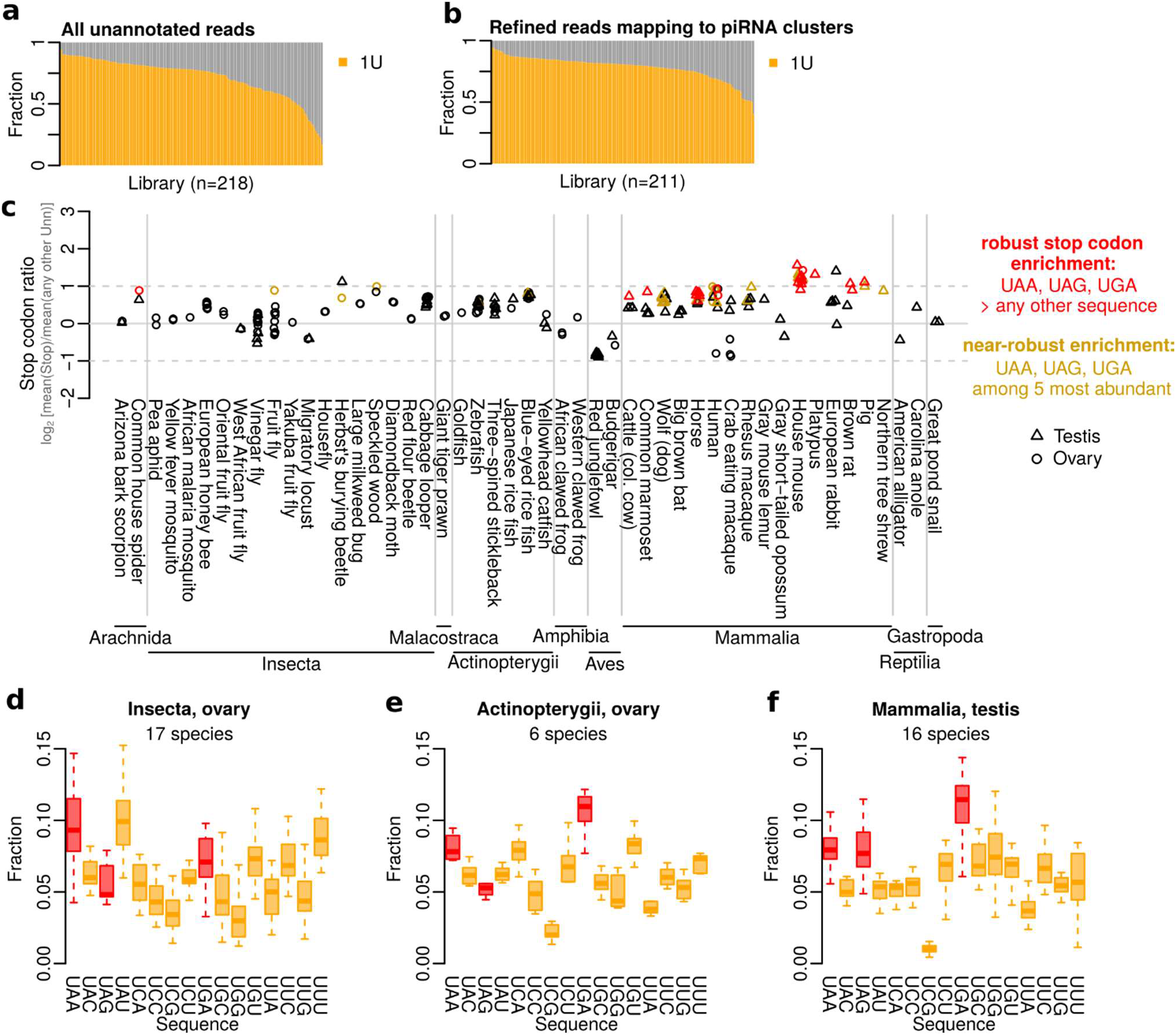
Stop codon enrichment is conserved across mammals. **a-b** Bar plots showing 1U across piCdb libraries. Showing either (a) all unannotated (i.e. piRNA) reads, or (b) unannotated reads mapping to piRNA clusters (see methods for details). **c** Stop codon ratio per testis (triangle) and ovary (circle) library calculated as mean frequency of stop codons vs mean frequency of all other U*nn* sequences. Signal shown on a log2 scale. Libraries with robust (red) or near-robust (yellow) stop codon enrichment are indicated. **d-f** Boxplots showing relative fraction of U*nn* sequences in insect ovary (d), ray-finned fish ovary (e), or mammalian testis (f). Each species is represented as one data point in the boxplots, and multiple libraries from the same species were averaged. The three largest groups are shown here, see also Fig. S9. Boxplots show median (central line), interquartile range (IQR, box), and minimum and maximum values (whiskers, at most 1.5*IQR).

To investigate the stop codon enrichment further, we identified libraries in which all three stop codons were found more frequently at the 5’ end than any other sequence (hereafter referred to as a *robust* stop codon enrichment). This revealed 23 libraries (11% of all libraries; expected fraction if any U*nn* was equally likely is 0.18%) with a robust stop codon enrichment (Fig. 6c) and an additional 26 libraries with a *near-robust* stop codon enrichment, where all stop codons were among the five most abundant sequences. The 23 libraries with a robust enrichment comprised 22 libraries from eight mammals (cattle [*Bos taurus*], horse [*Equus caballus*], common marmoset [*Callithrix jacchus*], human, mouse, pig [*Sus scrofa*], platypus [*Ornithorhynchus anatinus*], and brown rat [*Rattus norvegicus*]) and one library from arachnids (common house spider [*Parasteatoda tepidariorum*]). The libraries with near-robust stop codon enrichment (Fig. 6c) included another 21 libraries from mammals covering three additional mammalian species, two from ray-finned fish, and three from insects. Notably, libraries with robust stop codon enrichment were strongly over-represented in the mammalian group with 22 out of 80 (28%) of mammalian libraries against 1 out of 132 (0.8%) of remaining libraries (p=1e-9, Fisher’s exact test). Intriguingly, while the mammalian group included rat, a rodent that diverged from mouse only 21 million years ago (MYA), it also included the evolutionary much more distant species human and common marmoset (90 MYA), pig, horse, and cattle (96 MYA), and platypus (177 MYA), suggesting that a 5’ end stop codon enrichment is conserved across the entire mammalian lineage.

The analysis above was based on individual small RNA-seq libraries. To exclude the possibility that variability in the number of libraries per species affected the global analysis, we next averaged the trinucleotide frequencies for libraries from the same species and tissue (ovary or testis) to give each species equal weight. Supporting the previous observations, among all class and tissue combinations (Fig. 6d-f, Fig. S9a-h), the mammalian testis group that included 16 species displayed a robust stop codon enrichment (Fig. 6f). In addition, the mammalian ovary group showed a near-robust stop codon enrichment (Fig. S9g) with overall frequencies very similar to testis (Fig. S9i). We concluded that the stop codon enrichment at piRNA 5’ ends is conserved across mammals.

### Sequence composition of piRNA clusters does not explain stop codon enrichment

The three stop codons occur frequently in the genome and are only absent in-frame within coding regions. If all bases were equally likely, stop codons would occur in 3 out of every 64 positions. However, piRNA clusters often have a low GC content, meaning that stop codons may occur more frequently. We next asked whether the underlying trinucleotide frequencies in piRNA clusters contributed to either the presence or absence of a stop codon enrichment at piRNA 5’ ends across the 49 species. For this analysis, we used the predicted piRNA cluster transcripts in the piRNA cluster database ^43^, excluded cluster transcripts that were not expressed, and then divided them into all possible 3-mers. This empirical trinucleotide distribution was used to derive the expected relative distribution of U*nn* sequences. A normalized sequence composition was defined as the ratio between the previously observed piRNA 5’ end sequence distribution to the one expected based on the cluster composition, as illustrated in Fig. 7a.

**Fig. 7.**
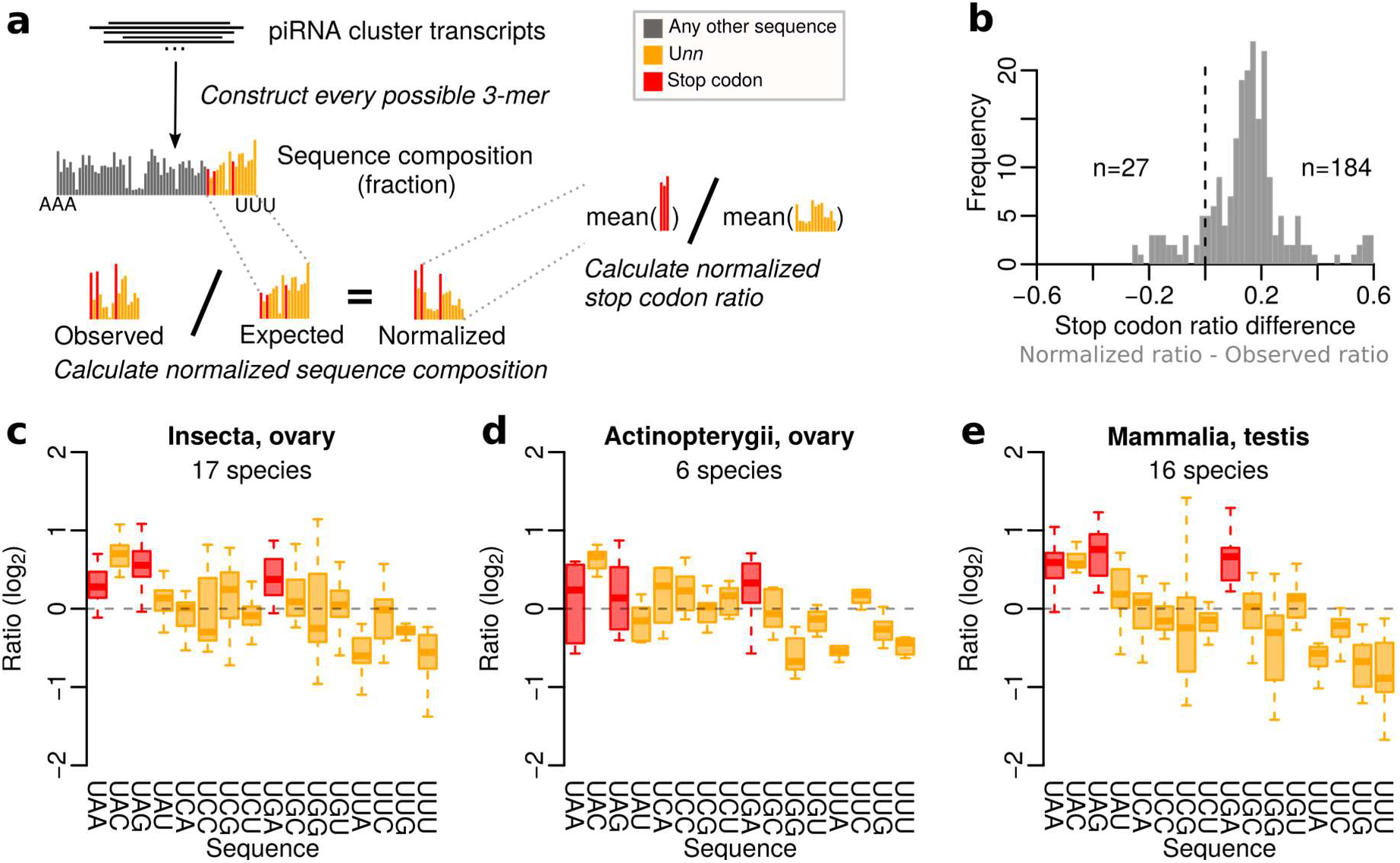
Stop codon enrichment is independent of piRNA cluster sequence composition. **a** Schematic to illustrate the data processing. First, piRNA cluster transcripts are used to construct the expected piRNA 5’ end sequence distribution. Second, the observed piRNA 5’ end sequence composition is divided by the expected sequence distribution to construct the normalized sequence composition. Third, stop codon ratio is calculated based on the normalized sequence composition. **b** Histogram showing the difference in stop codon ratio with or without normalization to the piRNA cluster composition. See also Fig. S10a-b for normalized ratios and differences per library. **c-e** Boxplots showing ratio between observed and expected fraction of U*nn* sequences in insect ovary (c), ray-finned fish ovary (d), or mammalian testis (e). Each species is represented as one data point in the boxplots, and multiple libraries from the same species were averaged. The three largest groups are shown here, see also Fig. S11. Boxplots show median (central line), interquartile range (IQR, box), and minimum and maximum values (whiskers, at most 1.5*IQR).

Surprisingly, this normalization increased the relative stop codon enrichment at piRNA 5’ ends across 184 out of 211 libraries (87%) representing the vast majority of species (Fig. 7b, Fig. S10a-b). An increased number of mammalian libraries displayed a robust or near-robust stop codon enrichment and similar patterns appeared across four ray-finned fish species and across insects such as moths and butterflies (Fig. S10a). Furthermore, by calculating the mean enrichment across all libraries from the same species and tissue (ovary or testis) and grouping the results by class and tissue (Fig. 7c-e, Fig. S11a-i), we confirmed that a stop codon enrichment was still present in the mammalian testis group (Fig. 7e). However, robust or near-robust stop codon enrichment was also found in the insect ovary (Fig. 7c), insect testis (Fig. S11c), amphibian ovary (Fig. S11e), mammalian ovary (Fig. S11g), and reptilian testis (Fig. S11h) groups. Notably, the UAC sequence were highly enriched in most groups, suggesting that the same four sequences are preferred cleavage positions across a wide range of species. We further noticed that libraries with poor 1U signal often have low stop codon ratio (Fig. S12, Fig. S13), suggesting that library quality or the presence of other processing such as ping-pong may influence the observed stop codon enrichment. Indeed, we observed a positive correlation between 1U and stop codon ratio both before (Fig. S10c, r^2^=0.20, p=9e-12) and after (Fig. S10d, r^2^=0.23, p=2e-13) normalizing the libraries to the underlying cluster composition.

Our analysis shows that stop codon enrichment at mammalian piRNA 5’ ends is independent of the piRNA cluster sequence composition. Moreover, although only one species outside of mammals exhibited a robust stop codon enrichment (Fig. 6c), there are nevertheless indications that piRNA biogenesis favours 5’ end stop codons relative to other sequences in many non-mammalian species. Thus, the mechanism underlying stop codon enrichment at piRNA 5’ ends may be shared wider across the animal kingdom.

## DISCUSSION

Here we identified and characterized a previously unknown enrichment of stop codons (UAA, UAG, UGA) at mouse piRNA 5’ ends, around 2-fold stronger than the well-known 1U bias. The 1U bias was noted when piRNAs were first described and is the hallmark of the Zuc-dependent phased biogenesis ^14,15^. Nevertheless, the source of the 1U bias is still unclear as the Zuc endonuclease is believed not to have any intrinsic sequence preference ^18,19^. Recently, an extended cleavage motif based on five additional positions with relative enrichments around 1.2-1.7-fold was described for silkmoth Zuc ^41^. By performing a similar analysis in mouse, we revealed a -1G depletion as the strongest extended motif, which was also partially present in silkmoth but not part of the reported motif ^41^. The second strongest position in mouse was a 5A signal, that was nearly 2-fold enriched in our pooled analysis, although absent in many individual replicates. The same position was reported as a C in the silkmoth Zuc motif, with an enrichment of 1.3-1.5-fold ^41^. Thus, sequences preferred by mouse PLD6 show high nucleotide variability and appear different to the silkmoth motif. In sharp contrast to the variable and weak enrichment of individual nucleotides, the enrichment of three distinct stop codon sequences was consistently present in all mouse libraries and in many other species. This indicates that at least in mammals, it is specific sequences— not individual nucleotides—that determine Zuc cleavage specificity *in vivo*. Further modelling and analysis indeed revealed a conserved cleavage preference at four sequences, corresponding to the three stop codons and the low-abundant UAC sequence, which specifically promotes piRNA 5’ end formation at stop codons.

While U occurs frequently in piRNA cluster transcripts, only some Us occur in a stop codon context and these are randomly distributed. Thus, a strong preference for piRNA precursor cleavage at Us in a stop codon context should result in a very wide pre-piRNA length distribution. Indeed, most animals have pre-piRNAs reaching 40 nt and sometimes up to 60 nt in length ^31^, whereas mature piRNAs rarely exceed 32 nt. Therefore, any cleavage preference beyond 1U would require efficient trimming of pre-piRNAs into mature piRNA size. To directly produce pre-piRNAs at mature piRNA length, no pronounced Zuc sequence specificity could be present. Notably, in a study that estimated pre-piRNA length and compared it to mature piRNA length across 35 species, only *D. melanogaster* displayed a pre-piRNA size distribution consistent with a nearly complete lack of trimming ^31^. We note that in agreement with this, we have been unable to detect any stop codon enrichment in *D. melanogaster*, despite the availability of large amounts of high-quality data. This supports the idea that any cleavage preference beyond 1U has likely been lost in *D. melanogaster*, and the remaining 1U bias may be what maintains the occasional but non-essential trimming of only a few nucleotides that has been previously described ^30^.

What is the function of a U or stop codon at piRNA 5’ ends? The first position of the guide piRNA is anchored in the 5’ phosphate-binding pocket within the PIWI protein and does not participate in base pairing with the target RNA ^44–47^. The 1U and the first position of a stop codon therefore appear to have limited influence on target recognition. It is likely that both 1U and stop codon enrichment have no immediate downstream function, but may instead be the signature of a yet unidentified process coupled to Zuc-dependent phased biogenesis. This is in line with the lack of purifying selection targeting the stop codon positions. Since stop codons were selectively enriched this naturally provokes speculation that this unknown process may include or have evolved from factors involved in translation, including RNA surveillance such a monitoring of codon optimality ^48^. We speculate that preferential targeting of stop codons may have evolved to tune the machinery toward non-coding piRNA precursors rather than coding sequences. Zuc-mediated cleavage occurs on the mitochondrial outer membrane, which is also the site of translation of many mRNAs. Ribosome profiling has revealed that nearly half of all mouse non-coding RNAs are associated with ribosomes ^49^, and piRNA precursors are likely candidates to be among them due to their cytoplasmic localization. Indeed, both piRNA precursors and PLD6 have been observed to associate with polysomes ^33^. Moreover, MIWI loaded with a piRNA is capable of associating with polysomes in an RNA-dependent manner ^50^. Assessing translation through ribosome profiling in testis is challenging since contamination by highly abundant small RNAs—including piRNAs—is to be expected. Nevertheless, ribosome profiling experiments in mouse piRNA knockout testis (*Mov10l1* knockout) have detected translated 5’ end-proximal ORFs in several piRNA precursors ^33^, indicating that the observed polysome association is at least sometimes driven by translation.

Translation and codon usage were recently implicated in small RNA biogenesis in worms (*Caenorhabditis elegans*), where CSR-1-associated 22G-RNAs complementary to mRNA coding sequences are produced in phase with ribosomes ^51^. In addition, the level of translation was inversely correlated to 22G-RNA abundance, suggesting that elongating ribosomes antagonize the production of 22G-RNAs ^51^. In plants (*Arabidopsis thaliana*), ribosome stalling occurs 12-13 nt upstream of several miRNA binding sites, mediated by the binding of the plant-specific dsRNA-binding protein SGS3, which triggers the production of downstream secondary phased siRNAs ^52^. Although the mechanisms differ in each case, the stop codon enrichment at piRNA 5’ ends suggests that also piRNA biogenesis may be connected to the translational machinery.

We note that although it is plausible that the identified stop codon enrichment indicates a connection to the translational machinery, an alternative model is that the detected sequence enrichments represent a previously hidden Zuc cleavage motif that only coincidentally promotes 5’ end stop codons. Notably, a hypothetical U-purine-purine motif would also be consistent with an enrichment of the three stop codons. However, this motif would also predict UGG to be enriched, which we have neither observed in mouse (Fig. 1b, Fig. 4c) nor in other mammals (Fig. 6f, Fig. 7e), and does not explain the apparent enrichment of UAC.

In conclusion, we here report a previously unrecognized enrichment of stop codons at piRNA 5’ ends. This was discovered in mouse piRNA populations, but appears to be conserved in mammals and potentially further, including partial evidence seen in ray-finned fish and some insects. The cause and function of this enrichment is still largely unknown, but our study contributes novel insights into Zuc-mediated piRNA biogenesis and provides a solid foundation to resolve these questions, which may eventually connect piRNA biogenesis with the translational machinery.

## METHODS

### Retrieval and processing of small RNA-seq libraries

Processed sRNA-seq libraries data in bed2 or similar format were downloaded from the GEO ^53^ if available and otherwise downloaded as raw data from the SRA ^54^. The reads were filtered in two ways to enrich for piRNAs. First, reads corresponding to other types of RNAs such as miRNA, tRNA and rRNAs were generally excluded. Second, only reads with a length consistent with piRNAs in the corresponding stage were kept. Detailed information on the processing of each dataset is provided below. Only libraries with at least 1,000 reads after filtering are shown in Fig. 1 and Fig. 2.

*Mov10l1* knockout and control libraries with six replicates per condition (GSM4160768-GSM4160779) and one library representing untreated adult mouse testis (GSM4160780) were downloaded in bed2 format from the GEO under accession GSE65786 ^33^. These reads already had adapters removed, rRNAs and miRNAs excluded and had been aligned to the *mm10* reference genome as described previously ^33^. Reads mapping to annotated tRNA or rRNA locations (retrieved from the UCSC genome browser), or the mitochondrial genome were removed for the 5’ end analysis. Only reads of length 26-32 nt were included in the analysis.

*Pnldc1* knockout and control libraries, single-end 75 nt, with each condition done in quadruplicates under accession PRJNA421205 ^31^ were downloaded from the SRA. The libraries were downsampled to 2 million reads per library to speed up the processing. Adapters were removed using Trim Galore! (v0.6.4, --stringency 6 --three_prime_clip_R1 3 --length 26 --max_length 32), to also remove three random nucleotides near the 3’ adapter and to retain only reads of length 26-32 nt. Next, the reads were aligned against the *mm10* reference genome using TopHat2 (v2.1.1, --max-multihits 1 --no-coverage-search, Gencode vM24 gene models) ^55^ to identify uniquely mapping reads.

*A-Myb* knockout (six replicates from 14.5 dpp and two from 17.5 dpp) and heterozygote (four replicates from 14.5 dpp and two from 17.5 dpp), *Miwi* knockout and heterozygote (four replicates per genotype from 14.5 dpp and two per genotype from 17.5 dpp), *Spo1l* knockout and heterozygote (one replicate per genotype), and wild-type libraries from 10.5, 12.5, 14.5, 17.5, 20.5 days and 6 weeks post partum (two replicates per time point) single-end 50 nt libraries were downloaded from the SRA under accession PRJNA194540 ^7^. The libraries were downsampled to 1 million reads per library to speed up the processing. Adapters were removed using Trim Galore! (v0.6.4, --stringency 6 --length 26 --max_length 32) to retain only reads of length 26-32 nt. Next, the reads were aligned against the *mm10* reference genome using TopHat2 (v2.1.1, --max-multihits 1 --no-coverage-search, Gencode vM24 gene models) to identify uniquely mapping reads.

*Mili* knockout, wild-type, MILI-IP in *Dnmt3l* knockout, and MILI-IP in wild-type libraries from 10 dpp and MILI-IP, MIWI2-IP, and total libraries from 16.5 dpc were downloaded from the SRA under accession PRJNA111011 ^34^. The libraries were single-end with read length 30-36 nt. Adapters were removed using Trim Galore! (v0.6.4, --stringency 3 --length 26 --max_length 32 -a CTGTAGGCACCATCAATCGTATGCCGTCTTCTGCT) to retain only reads of length 26-32 nt. Prenatal libraries instead used --length 23 to retain reads of length 23-32. Next, the reads were aligned against the *mm10* reference genome using TopHat2 (v2.1.1, --max-multihits 20 --no-coverage-search, Gencode vM24 gene models) followed by extraction of primary alignments using samtools. Reads corresponding to the snRNA *Snord2* (CTGAAATGAAGAGAATACTCTTGCTGATC) were excluded from the 5’ end analysis.

*Tex15* knockout and heterozygote control libraries with three replicates per genotype and corresponding MILI-IP and MIWI2-IP libraries from with two replicates per genotype (GSM2881231-2881244) were downloaded in processed format from the GEO under accession GSE107832 ^37^. These reads already had adapters removed, and had been annotated and aligned to the *mm10* reference genome as described previously ^37^. Reads previously identified as miRNAs or ncRNAs and reads overlapping annotated tRNA or rRNA locations (retrieved from the UCSC genome browser) or the mitochondrial genome were excluded. Only reads of length 23-32 nt were included in the analysis.

*Tex15* knockout and heterozygote control libraries with three replicates per condition (GSM4547340-4547345) were downloaded in processed format from the GEO under accession GSE150350 ^38^. These reads already had adapters removed and had been annotated as described previously ^38^. Reads previously identified as miRNAs or ncRNAs were excluded from the analysis. Only reads of length 23-32 nt were included in the analysis and we counted at most 100 copies of the same sequence to remove abundant contaminants.

*Mili* and *Pld6* knockout and control libraries with three replicates per condition, *Dicer* knockout and a heterozygote control, *Miwi* and *Nanos3* knockouts, and oxidized and non-oxidized libraries wild-type (one replicate each) from 20 dpp ovaries, as well as MILI-IP in *Mili* knockout or control ovaries (one replicate each) from 10 dpp ovaries (DRR083999-DRR084018) were downloaded from the SRA under accession PRJDB4628 ^36^. These reads already had adapters removed and were of length 18-52. The reads were aligned against the *mm10* reference genome using TopHat2 (v2.1.1, --g 1 --no-coverage-search, Gencode vM24 gene models). Reads mapping to annotated tRNA or rRNA locations (retrieved from the UCSC genome browser), or the mitochondrial genome were removed for the 5’ end analysis. Only reads of length 26-32 nt were included in the analysis and we counted at most 50 copies of the same sequence to reduce the effect of abundant contaminants.

*Pld6* knockout and control libraries from 16.5 dpc and *Pld6* knockout from 10 dpp with one replicate per condition (GSM509275, GSM509276, GSM509280) were downloaded the GEO under accession GSE20327 ^35^. All libraries had read length 36 nt. Adapters were removed using Trim Galore! (v0.6.4, --stringency 3 -q 0 –length 20 --max_length 35 -a CGTCGTATGCCGTCTTCTGCTTGT) to retain only reads where adapters could be detected. Next, the reads were aligned against the *mm10* reference genome using TopHat2 (v2.1.1, --g 1 --no-coverage-search, Gencode vM24 gene models). Reads mapping to annotated tRNA or rRNA locations (retrieved from the UCSC genome browser), or the mitochondrial genome were removed for the 5’ end analysis. Only reads of length 26-32 nt (postnatal) or 23-32 nt (prenatal) were included in the analysis and we counted at most 50 copies of the same sequence to reduce the effect of abundant contaminants.

### Additional processing of Pnldc1 knockout and control small RNA-seq libraries

For the *Pnldc1* knockout and control libraries that were used outside of the analysis in Fig. 1 we also processed the libraries to include all reads with no down-sampling or size selection. For this analysis, adapters were removed using Trim Galore! (v0.6.4, --stringency 6 -q 0 --three_prime_clip_R1 3 --max_length 68), followed by alignment against the *mm10* reference genome using TopHat2 (v2.1.1, -g 1 --no-coverage-search, Gencode vM24 gene models). This strategy retained uniquely mapping reads of all sizes with known 5’ and 3’ ends.

### Determination of piRNA 3’ end downstream sequence

This analysis used the mouse *Pnldc1* knockout and control libraries described above to extract the nucleotide sequence immediate downstream of piRNA 3’ ends. Reads with alignment mismatches were excluded to avoid non-templated piRNA tailing. BEDOPS was used to convert bam files to bed, reads mapping to chrM were removed, bedtools intersect was used to exclude reads overlapping tRNA, rRNA or miRNA, and to restrict the analysis to reads mapping to pachytene piRNA clusters. Flanking sequence immediately downstream of the reads was extracted using bedtools flank followed by getfasta in strand-specific mode.

### Mouse pachytene piRNA cluster annotations

For mouse pachytene piRNA clusters we used the clusters or the “uORF downstream region” or the full cluster coordinates previously reported ^33^.

### piRNA 5’ end definition score

The purpose of this analysis was to study how the local sequence composition within piRNA precursor transcripts affected the likelihood for cleavage to happen. In short, we scored each transcript position by the observed cleavage likelihood and compared scores across different nucleotide and trinucleotide contexts. The 5’ end definition score was calculated using six control libraries from adult mouse testis ^33^. We used exonic regions in the mouse pachytene piRNA cluster uORF downstream region as the regions of interest.

To describe the competition between potential cleavage sites, we considered positions within 10 nucleotides upstream and 20 nucleotides downstream as possible alternative cleavage sites. For each position *x* within a cluster, the 5’ end definition score, *score*5, was defined as

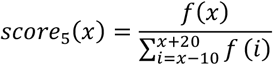

where *f*(*x*) is the number of 5’ ends mapping at position *x*. This score reflects the fraction of 5’ ends that map exactly to position *x* within this local region. To focus on high-confidence scores, the subsequent analyses of the scores were limited to positions with ≥100 reads within the local region.

In contrast to similar modelling that has been done for piRNA 3’ ends previously ^30^, our score does not require reducing the data to regions with a single strong phasing pattern but can be calculated across any region with sufficient read coverage.

Similar 5’ end definition scores were also calculated using sorted cells from the *Pnldc1* knockout (n=24) and control (n=28) libraries ^31^. This was done using both the observed 5’ ends (at the 5’ ends of the reads) and the expected ones assuming a phased processing (immediately downstream of 3’ ends). To make the different datasets more comparable, we used the score thresholds derived from the initial analysis.

### Identification of stop codons in ORFs

To investigate whether the presence of an ORF affected the likelihood for a stop codon to become a cleavage site, we identified 92,481 stop codons present in the mouse pachytene piRNA precursor uORF downstream regions. Out of these, 25,093 were part of an ORF and 67,388 were not. Coordinates for the stop codons were intersected with the piRNA 5’ end definition scores (described above) to exclude regions with low read coverage (<100 reads within the local region) and to assign a score to each stop codon. Samples from the control condition were used for this analysis (6 replicates) and stop codons covered by multiple replicate were assigned one score for each replicate. In total, this gave 94,472 data points, representing 22,374 stop codon positions that were part of an ORF of length 3-100 aa and 59,227 positions there were not. They were further subdivided into three groups of similar size (3-8 aa, 9-19 aa, and 20-100 aa), to test if ORFs of a particular size were associated with piRNA 5’ ends. Differences were assessed through Wilcoxon rank sum test.

To analyse experimentally confirmed ORFs, we used a set of 19,023 ORFs with experimental evidence from ribosome profiling and/or proteomics ^42^. This identified 28 ORFs within pachytene piRNA clusters. All 28 ORFs had transcriptional evidence in testis and 19 also had translational evidence. These regions were intersected with the piRNA 5’ end definitions scores to focus on six regions with sufficient read coverage. To assess whether their mean 5’ end definition score different from the expected one, we determine the distribution of the mean scores of control regions. The selection of control regions was made to mimic the observed ORF size distribution and was repeated 10000 times. The empirical p-value was determined as a fraction describing how often the observed score was lower than that of the control regions.

### Sequence conservation analysis using phyloP scores

To determine the sequence conservation across individual positions in mouse pachytene piRNA clusters we downloaded phyloP scores, representing a multiple alignment of 59 vertebrate species to the mouse genome, from the UCSC genome browser. We used phyloP scores both from the placental (40 species) and glire (8 species) subsets. The phyloP conservation scores were intersected with the piRNA 5’ end definition scores, to allow for comparisons in conservation between different 5’ end definition score bins. Additionally, bins were subsetted to only include U or stop codon positions in the comparisons. Conservation scores were represented by their mean score and the 95% confidence interval of the mean. A position was considered to show signs of positive selection if the associated phyloP score was negative and significant (i.e., phyloP < −1.30103). To assess sequence variation within mouse we used variants annotated in Ensembl r101 corresponding to dbSNP build 150. Annotated SNPs were intersected with the piRNA 5’ end definition scores and analysis were done per piRNA 5’ end definition bin, comparing the overall frequency of annotated variants for each bin, with or without restricting the analysis only to positions with a U.

### Conservation analysis using the piRNA cluster database

The piRNA cluster database ^43^ was used to extend the analysis in an unbiased way across 211 testis and ovary libraries from 49 species. All libraries can be interactively explored through the piRNA cluster database web interface (www.smallrnagroup.uni-mainz.de/piRNAclusterDB). This includes number of reads, read composition, positional nucleotide composition, length distribution, and pong-pong signature.

Scripts developed to retrieve and process all libraries have been made available on GitHub (www.github.com/susbo/5prime_stop_paper). In short, predicted piRNA cluster coordinates and sequence FASTAs were downloaded for all 51 species covered in the piRNA cluster database (a full list of species is available in Supplementary Table S1). Missing FASTA files from 12 species were re-created using the cluster coordinates and the listed reference assemblies. Focusing on all libraries derived from testis or ovary, we downloaded small RNAs with “unknown” annotation (i.e., putative piRNAs) from the database. In total, 225 libraries from 49 species were included in our study, after excluding five libraries that were either erroneously annotated as the wrong species (SRR6662680, SRR6662664) or that still had adapters attached to the reads (SRR578908, SRR578909, SRR363983).

To identify piRNA clusters that are expressed in testis and/or ovary, we mapped the “unknown” small RNAs from each library back onto the predicted piRNA clusters from the corresponding species using Bowtie (v1.2.3, -y -f -M 1 --best --strata -S -p 20 --chunkmbs 2000 --nomaqround). Clusters were considered to be expressed piRNA clusters if they had ≥100 uniquely mapped reads, ≥80% of the reads derived from the preferred strand, and ≥40% 1U. Next, to refine each library further, we extracted the subset of reads of length 24-31 nt mapping to the expressed piRNA clusters. To reduce the effect of a few extremely abundant contaminating sequences, we counted at most 100 sequences (selected at random) originating from each position in the clusters. However, the reported results were consistent across different subsampling thresholds (Fig. S14). Libraries were included in subsequent refined analysis if they had ≥10000 reads mapping to expressed piRNA clusters (n=211), or in the global analysis if they had ≥100,000 reads in total (n=218). The number of libraries per species is detailed in Supplementary Table S1.

To estimate the expected codon composition of piRNA clusters per library, we constructed sequence FASTAs covering the expressed piRNA clusters based on each individual library. Each set of piRNA clusters was divided into all possible 3-mers based on their sense strand nucleotide sequence, and the overall frequency of each codon was determined. To normalize the observed piRNA 5’ end codon composition by the underlying codon composition within piRNA clusters, we calculated the ratio between the observed divided by the expected distribution of codon starting with a U.

### Estimation of time of divergence

All pairwise divergence times shown are the estimated time in TimeTree ^56^.

## DATA AVAILABILITY

All sequencing data used in this study are publicly available. Mouse datasets are available through GEO (accession GSE20327, GSE65786, GSE107832, GSE150350) or SRA (accession PRJNA421205, PRJNA194540, PRJNA111011, PRJDB4628). Processed libraries from other species are available from the piRNA cluster database (http://www.smallrnagroup.unimainz.de/piRNAclusterDB).

## CODE AVAILABILITY

The scripts used to retrieve and process all libraries in the piRNA cluster database are available on GitHub (www.github.com/susbo/5prime_stop_paper). All data processing is described in this paper.

## ACKNOWLEDGEMENTS

We thank Hannon group members for useful discussions and in particular Evelyn L Eastwood for valuable comments on the manuscript. We thank Ildar Gainetdinov for providing feedback on the manuscript. We thank the Scientific Computing core facility at CRUK Cambridge Institute for HPC resources. This work was funded in whole or in part by Cancer Research UK (A21143) and the Wellcome Trust (110161/Z/15/Z). For the purpose of open access, the author has applied a CC BY public copyright license to and Author Accepted Manuscript version arising from this submission. GJH is a Royal Society Wolfson Research Professor (RSRP\R\200001).

## AUTHOR CONTRIBUTIONS

SB conceived the study, analysed and interpreted the data and prepared the figures. SB designed the experiments and wrote the manuscript with significant input from BC and GJH. BC and GJH supervised the study and obtained funding. All authors read and approved the final manuscript.

