## Supplementary Information for "An evolutionarily conserved stop codon enrichment at the 5’ ends of mammalian piRNAs"

**This PDF includes:**

Fig. S1. Postnatal testis piRNAs are enriched for stop codons at their 5' ends. Related to Fig. 1.
Fig. S2. Postnatal Zuc-dependent ovarian piRNAs are enriched for stop codons at their 5' ends.
Fig. S3. Size-distribution of pre- and perinatal testis libraries. Related to Fig. 2.
Fig. S4. Related to Fig. 3.
Fig. S5. Related to Fig. 4.
Fig. S6. Sequences downstream of pre-piRNA 3' ends are enriched for 1U and stop codons in *Pnldc1*-KO mice.
Fig. S7. High-scoring bins are enriched for 1U and stop codons in *Pnldc1*-KO mice.
Fig. S8. Open reading frames do not contribute to piRNA 5' end definition.
Fig. S9. Related to Fig. 6.
Fig. S10. Related to Fig. 7.
Fig. S11. Related to Fig. 7.
Fig. S12. Overview of 114 libraries from testis. Related to Fig. 6 and Fig. 7.
Fig. S13. Overview of 97 libraries from ovary. Related to Fig. 6 and Fig. 7.
Fig. S14. Stop codon enrichment can be detected using different subsampling thresholds.

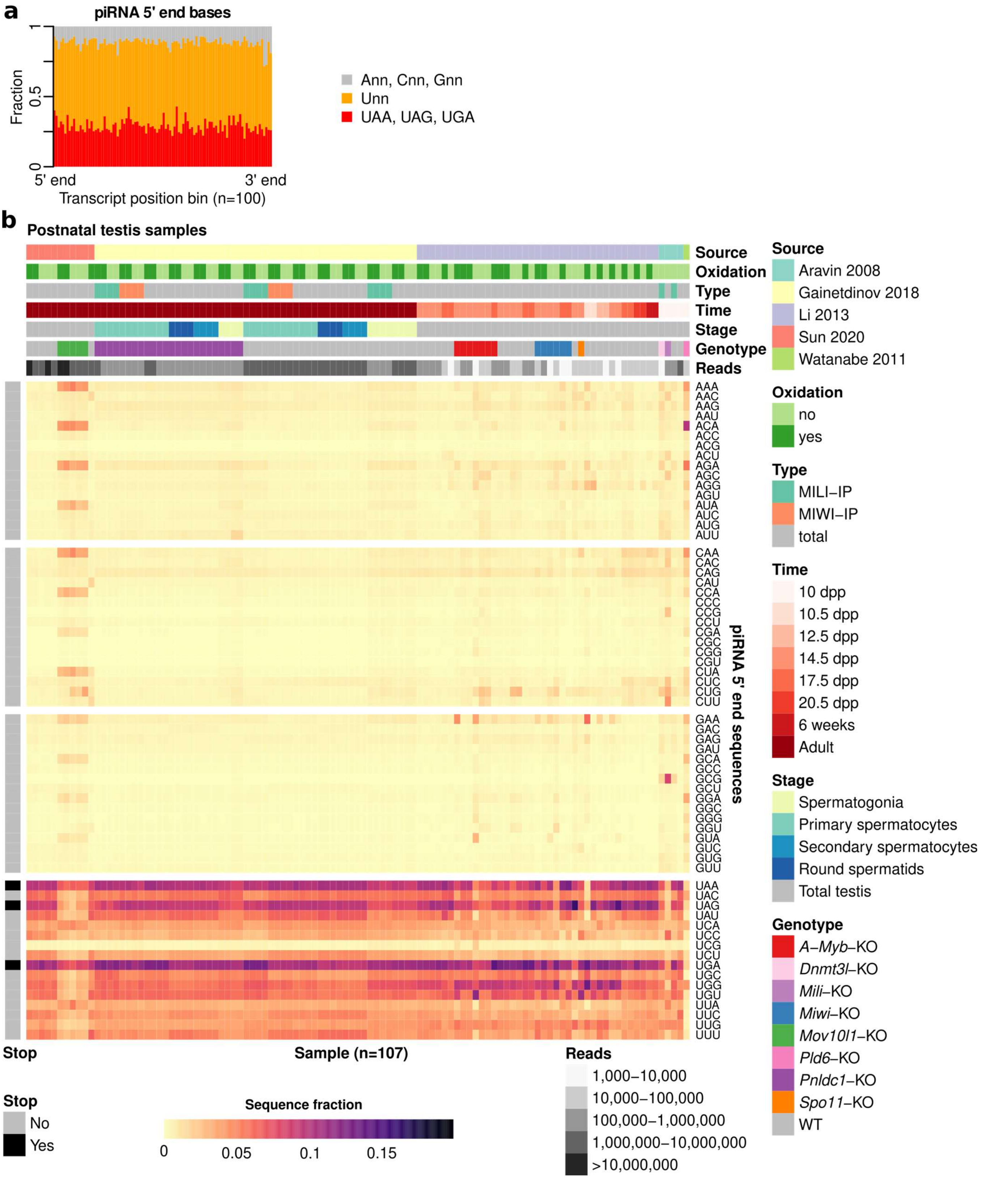


**Fig. S1. Postnatal testis piRNAs are enriched for stop codons at their 5' ends. Related to Fig. 1.**
**a** Mean 1U and stop codon fraction at piRNA 5' ends mapping to piRNA precursor transcripts. The precursor transcripts were divided into 100 equally sized bins, and the piRNAs mapping to each bin are shown separately. Pooled frequency for 5 adult wild-type replicates. **b** Heatmap showing 5' end sequence distribution of piRNAs. Rows display relative sequence distribution and columns represent 107 sRNA-seq libraries derived from postnatal mouse testis. Column-wise annotations describe data source publication (Source), whether oxidated RNAs were captured (Oxidation), library type (Type), developmental time point (Time), spermatogenesis stage (Stage), mouse genotype (Genotype), and number of reads (Reads). Row-wise annotations show whether a sequence is a stop codon (Stop). Abbreviations: dpp, days post partum; KO, knockout; WT, wild-type.

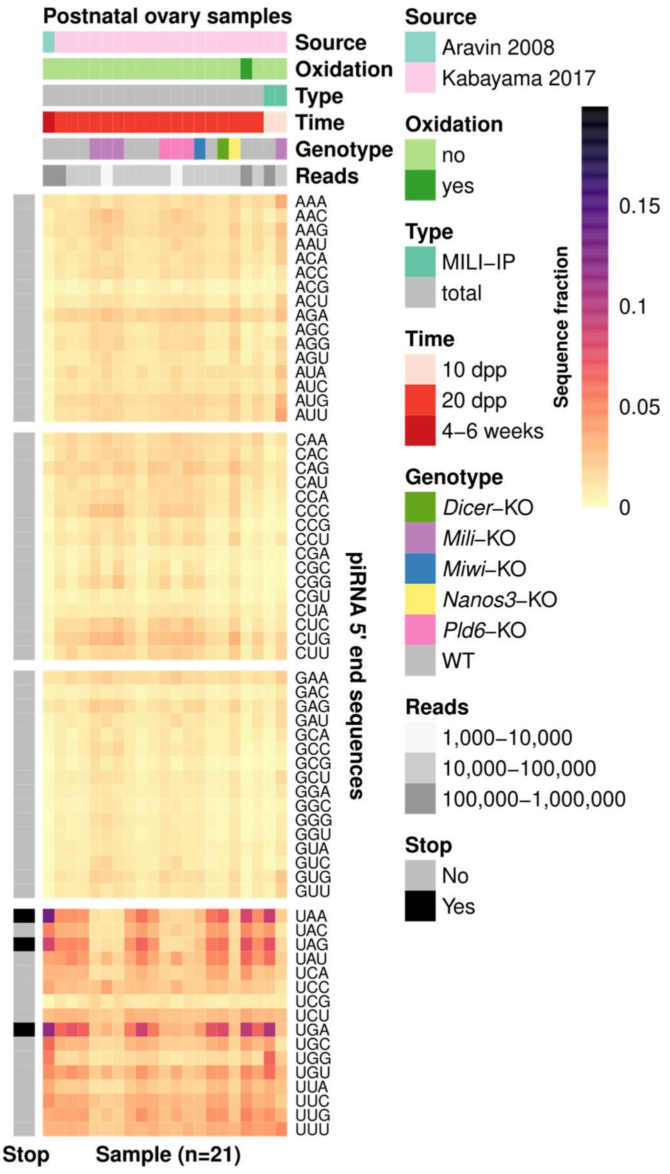


**Fig. S2. Postnatal Zuc-dependent ovarian piRNAs are enriched for stop codons at their 5' ends.**
Heatmap showing 5' end sequence distribution of piRNAs. Rows display relative sequence distribution and columns represent 21 sRNA-seq libraries derived from postnatal mouse ovary. Column-wise annotations describe data source publication (Source), whether oxidated RNAs were captured (Oxidation), library type (Type), developmental time point (Time), mouse genotype (Genotype), and number of reads (Reads). Row-wise annotations show whether a sequence is a stop codon (Stop). Abbreviations: dpp, days post partum; KO, knockout; WT, wild-type.

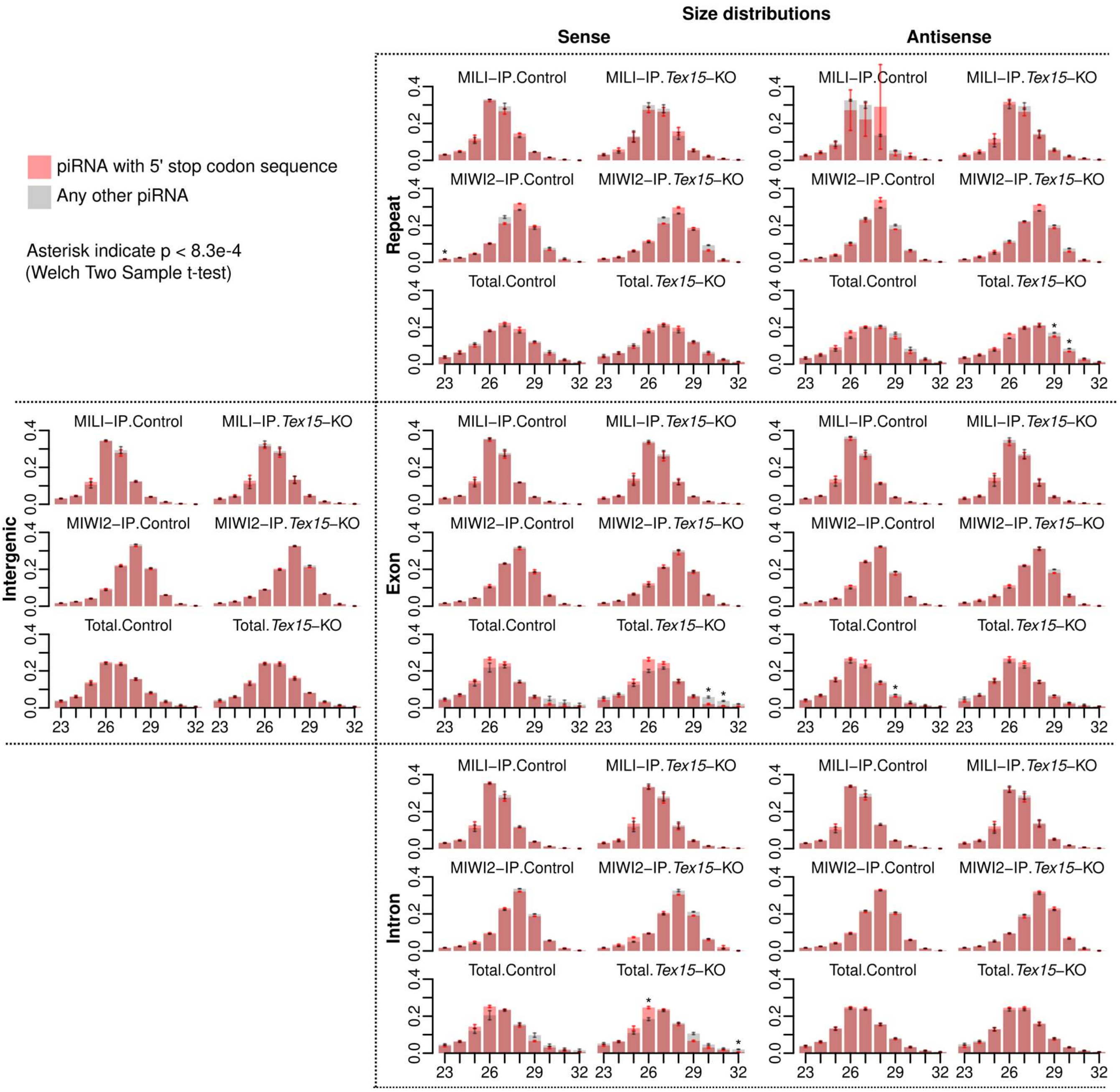


**Fig. S3. Size-distribution of pre- and perinatal testis libraries. Related to Fig. 2.**
Size distribution per annotation and strand for piRNAs with a stop codon sequence at their 5’ end (red) or all remaining piRNAs (grey). Bars represent mean fraction ± one standard deviation (2 replicates in IP libraries, 3 replicates in Total libraries). Difference between the two distributions was assessed per position using Welch Two Sample t-test and correction for multiple testing (Bonferroni correction with 60 tests).

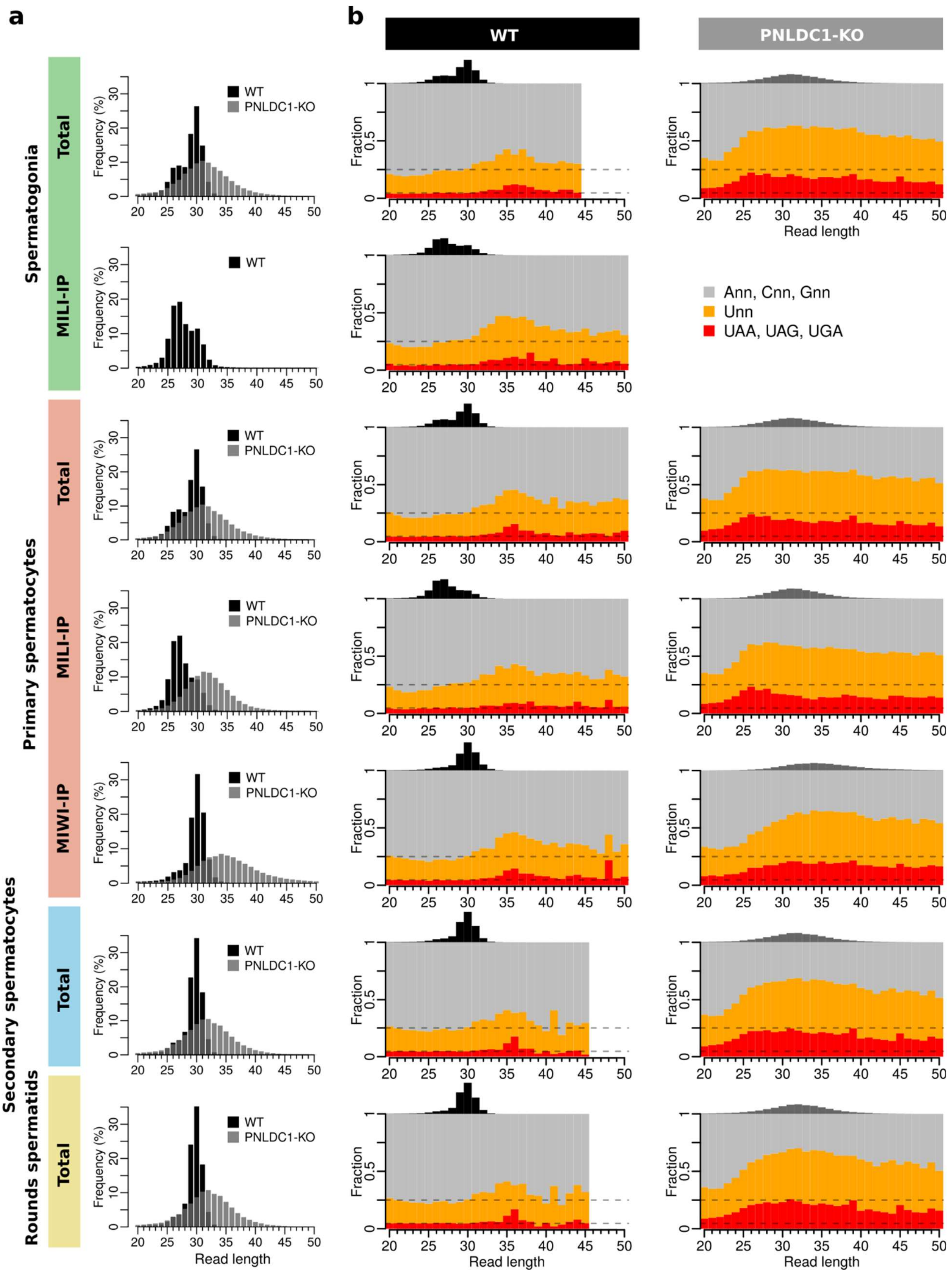


**Fig. S4. Related to Fig. 3.**
**a** Length distribution of PNLDC1 knockout (KO) (grey) and wild-type controls (black). Frequency shown as mean across four replicates. **b** 1U and stop codon fraction immediately downstream of 3' ends in wild-type (WT) and PNLDC1 knockout (KO) sRNA-seq libraries per read length. Dashed lines indicate expected fraction assuming all trinucleotide sequences are equally abundant. Fractions shown are mean across four replicates. Read lengths with <100 piRNAs are not shown.

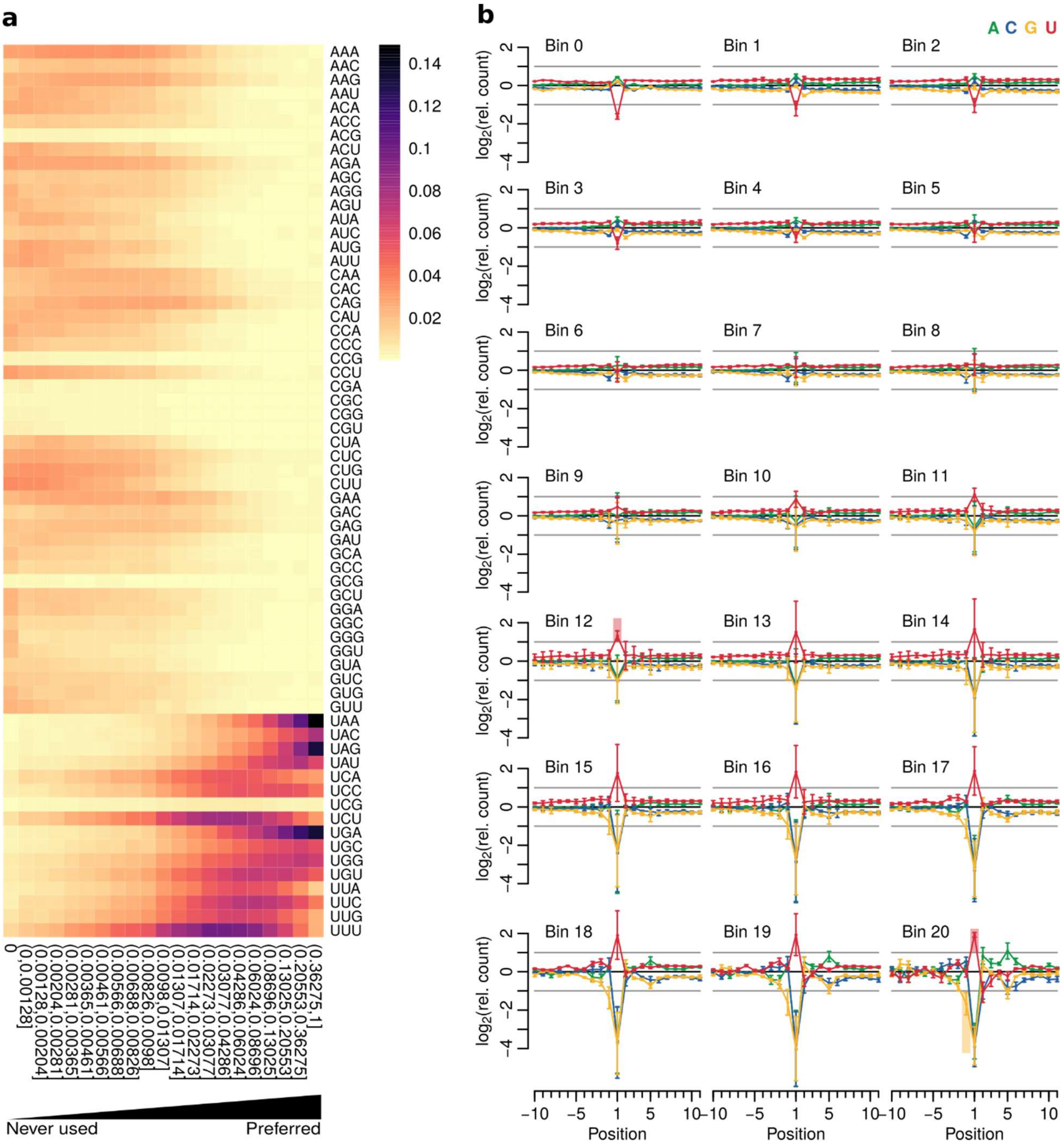


**Fig. S5. Related to Fig. 4.**

**a** Heatmap showing 5' end sequence distribution per bin. The 5' end definition score thresholds are shown under each column. Fraction calculated as mean across 5 replicates. **b** Nucleotide enrichment per position. Position 1 represent the piRNA 5' end and position -1 the position immediately upstream of the 5' end. Enrichment was calculated separately for each bin (pooled signal ± one standard deviation, 5 replicates). Bin 0 represent the least preferred positions and bin 20 to the most preferred ones.

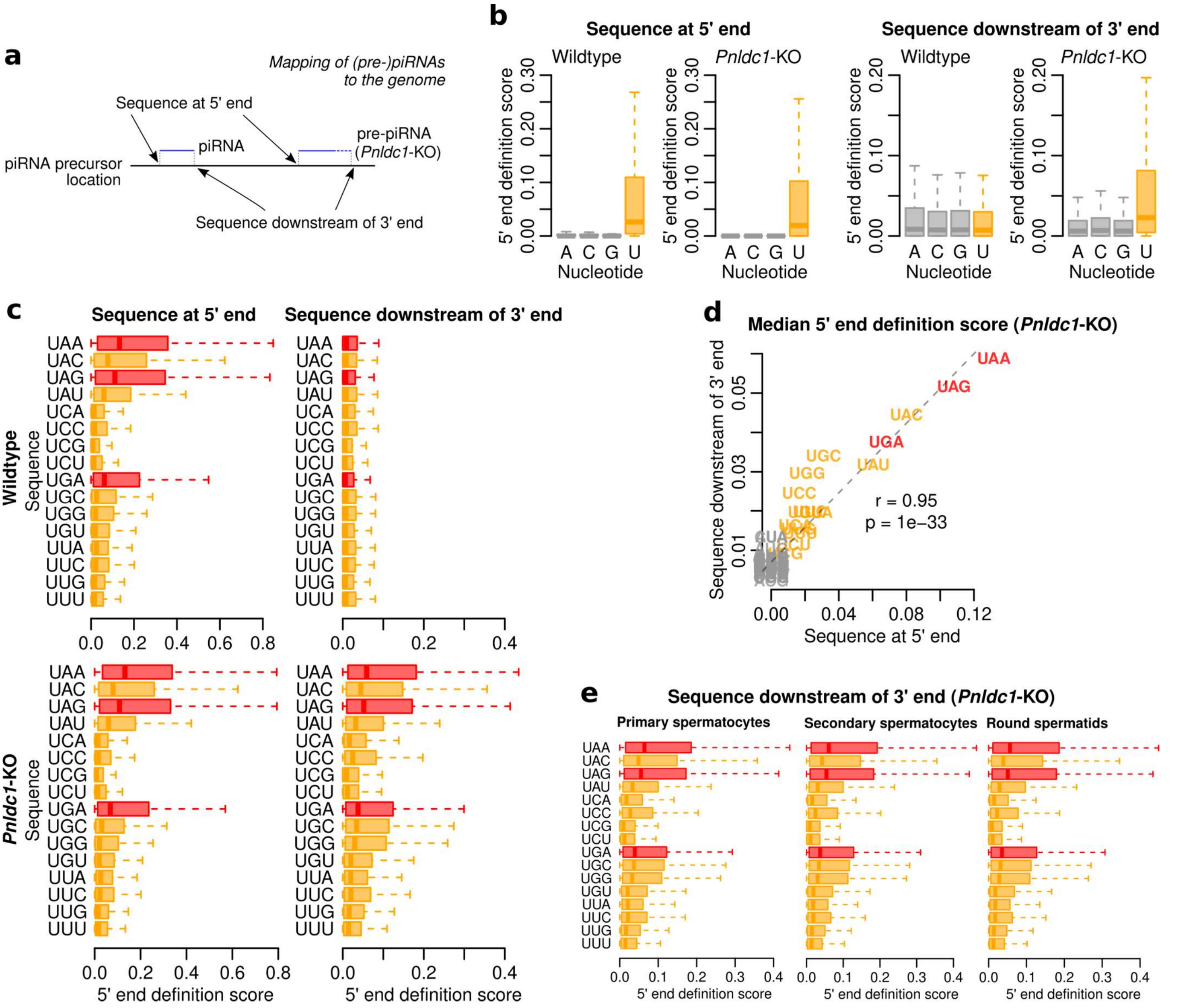


**Fig. S6. Sequences downstream of pre-piRNA 3' ends are enriched for 1U and stop codons in *Pnldc1*-KO mice.** **a** Illustration of sequences at (pre-)piRNA 5' end and downstream of 3' ends. *Pnldc1*-KO mice have defective piRNA 3' end trimming and those libraries will therefore capture longer piRNA intermediates (pre-piRNAs, extension indicated with a dashed line), whose 3' ends are expected to be positioned immediately upstream of the next piRNA. **b** Boxplot showing 5' end definition score per nucleotide using pooled data (28 wildtype or 24 *Pnldc1*-KO replicates). Sequences are derived either from piRNA 5' ends (left) or downstream of piRNA 3' ends (right). **c** Boxplot showing 5' end definition score per U*nn* sequence for wildtype (top, 28 pooled replicates) or *Pnldc1*-KO (bottom, 24 pooled replicates) mice using sequences from 5' ends (left) or 3' ends (right). Stop codons are shown in red and the remaining sequences in orange. **d** Scatterplot showing median 5' end definition score for sequences at 5' ends or downstream of 3' ends in *Pnldc1*-KO mice. Similarity between the medians was assessed using linear regression (dashed line). Pearson correlation and significance are indicated. **e** Same as (c) but separated on cell type as indicated. Pooled data (4 replicates each).

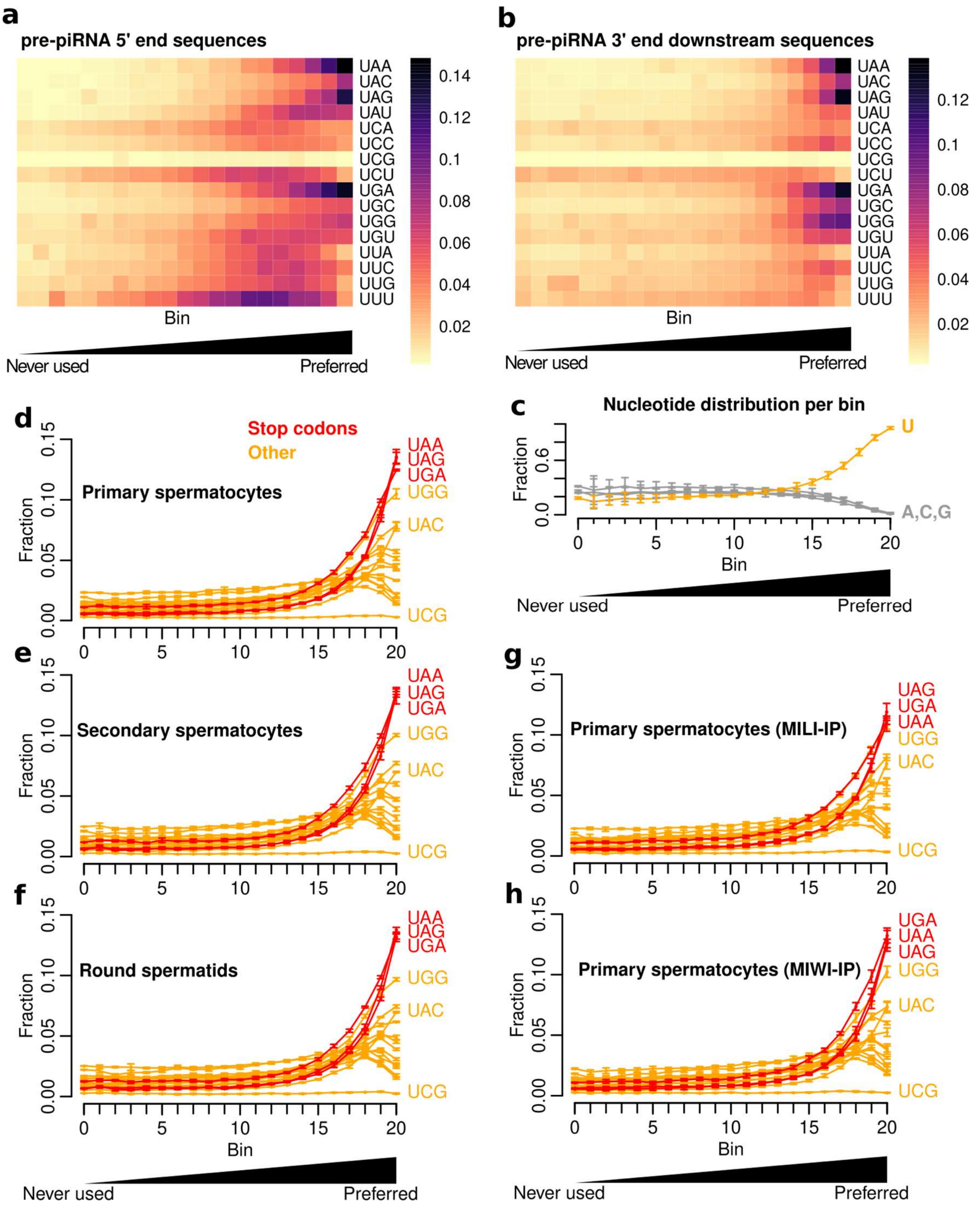


**Fig. S7. High-scoring bins are enriched for 1U and stop codons in *Pnldc1*-KO mice.**
**a-b** Heatmap showing sequence distribution at pre-piRNA 5' ends (a) or downstream of their 3' ends (b) per 5' end definition score bin. Binning thresholds were applied following Fig. 4f and reflect the least (bin 0) to the most (bin 20) preferred positions. Fraction calculated as mean across 24 *Pnldc1*-KO replicates. **c** Line graph showing nucleotide fraction per bin (defined as above) in *Pnldc1*-KO libraries. Fraction shown as mean ±sd (24 replicates). **d-f** Line graph showing U*nn* sequence fraction per bin in primary spermatocytes (d), secondary spermatocytes (e), or round spermatids (f) from *Pnldc1*-KO mice. Fraction shown as mean ±sd (4 replicates). **g-h** Line graph showing U*nn* sequence fraction per bin in MILI (g) or MIWI (h) immunoprecipitated libraries from primary spermatocytes in *Pnldc1*-KO mice. Fraction shown as mean ±sd (4 replicates).

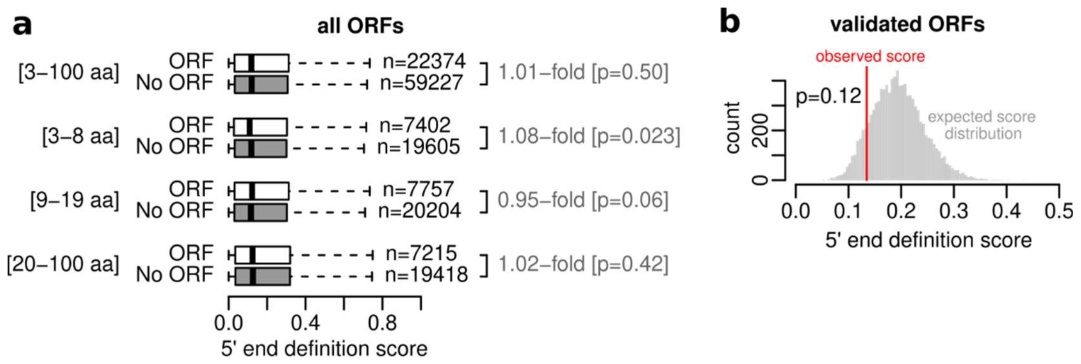


**Fig. S8. Open reading frames do not contribute to piRNA 5' end definition.**
**a** Boxplots showing 5' end definition score for stop codons either within (ORF) or outside (No ORF) of ORF context. The number of sites considered is indicated by the *n* value. The top row uses all ORFs and controls of 3-100 amino acids (aa) length. The three bottom rows explore using subsets of the data with different size ranges. Significance was accessed using Wilcoxon rank sum tests. Note that none of the presented p-values for the subset analysis is significant after correcting for multiple testing (i.e., Bonferroni correction) and the effect sizes are very minor. The observed minor differences are therefore not likely to represent a true biological difference. **b** Histogram comparing observed and expected 5' end definition score. The observed value (red line) is based on stop codons in ORF context validated by ribosome profiling, whereas the expected (grey histogram) is based on simulations selecting matched stop codons outside of ORF context. Empirical one-sided p-value.

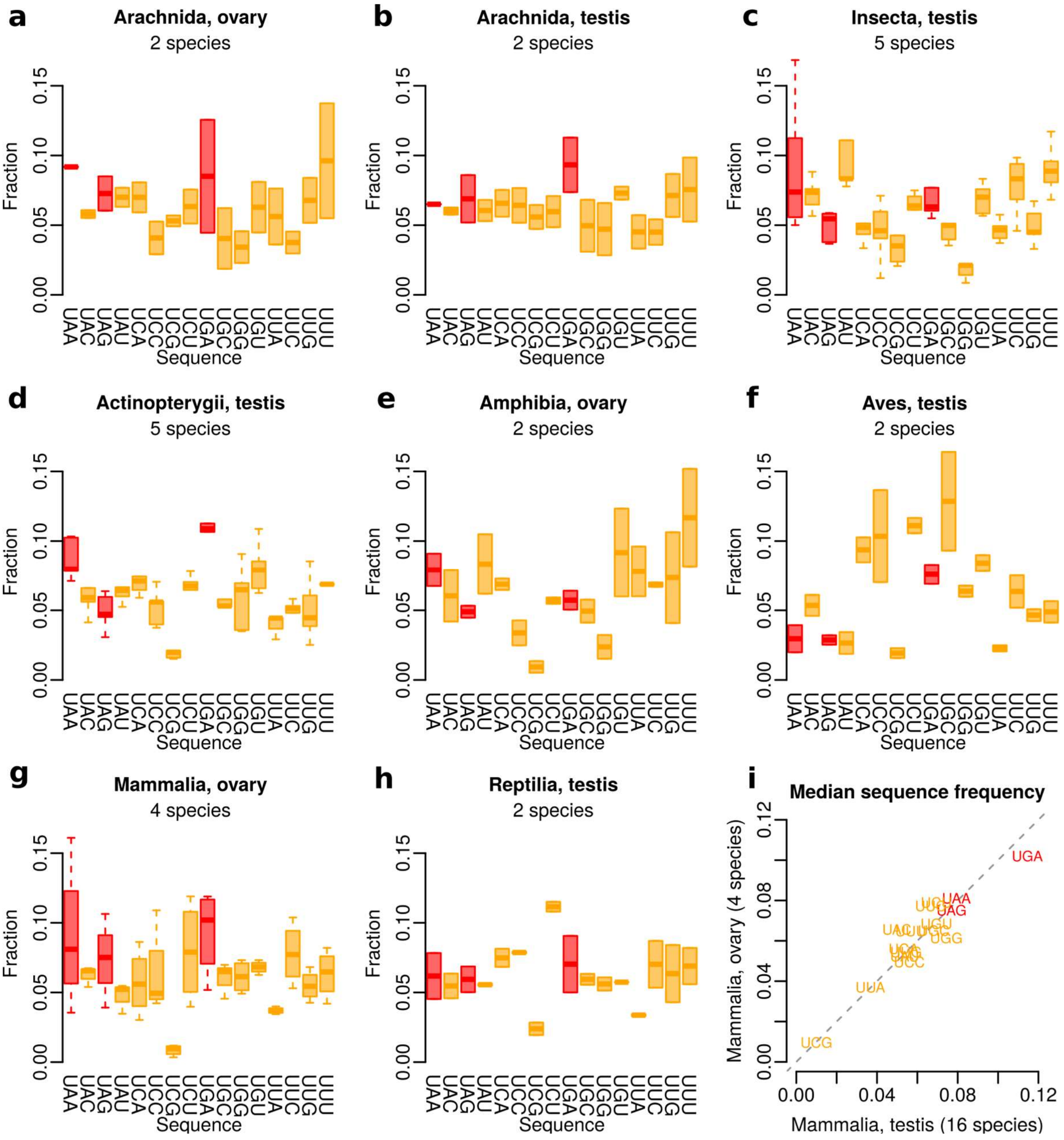


**Fig. S9. Related to Fig. 6.**

**a-h** Boxplots showing relative fraction of U*nn* sequences in arachnid ovary (a), arachnid testis (b), insect testis (c), ray-finned fish testis (d), amphibian ovary (e), bird testis (f), mammalian ovary (g), and reptilian testis (h). Each species is represented as one data point in the boxplots, and multiple libraries from the same species were averaged. Boxplots show median (central line), interquartile range (IQR, box), and minimum and maximum values (whiskers, at most 1.5*IQR). **i** Scatterplot comparing the median sequence frequency across mammalian species with ovary (Fig. S9g) and testis (Fig. 6f) libraries.

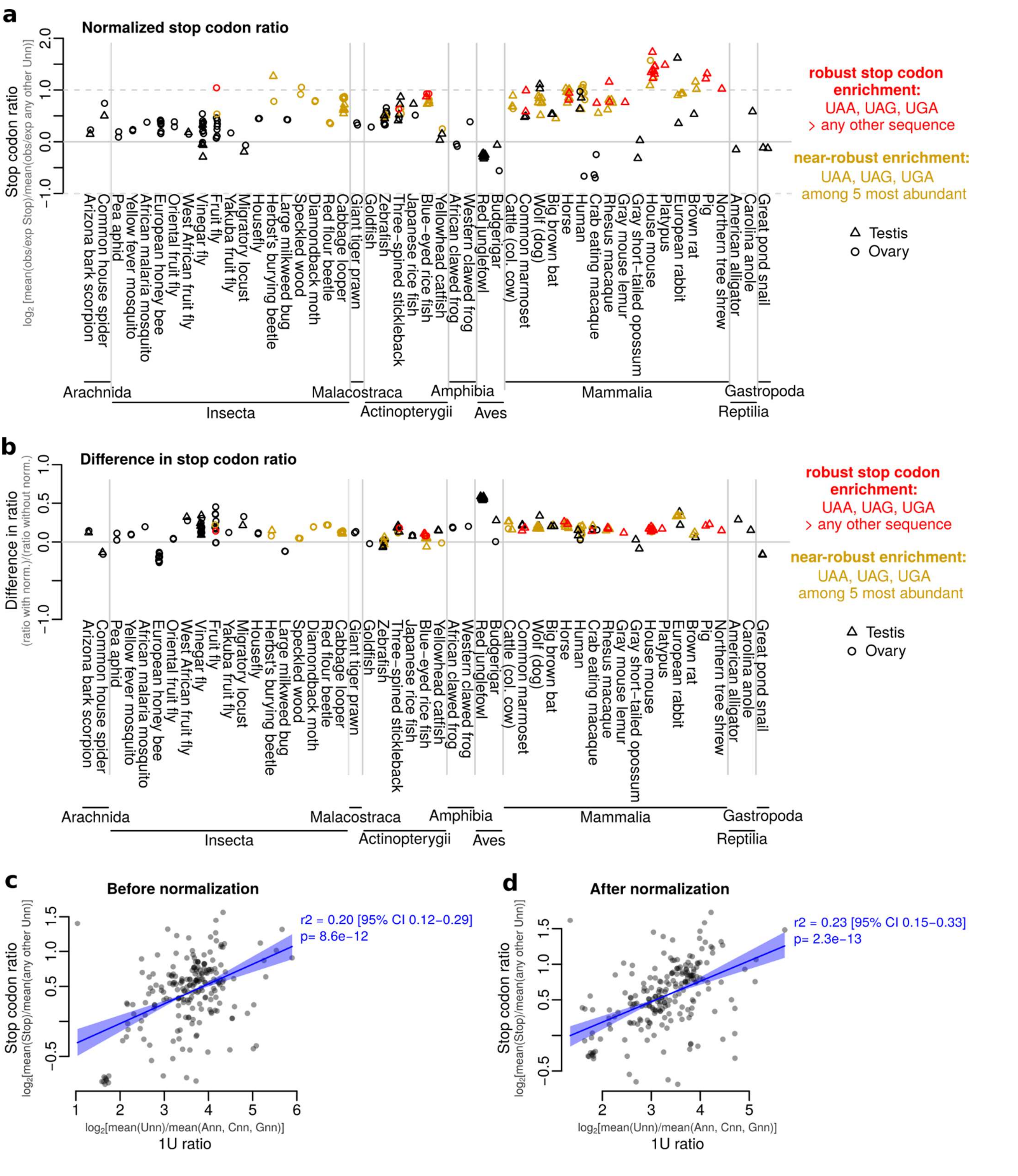


**Fig. S10. Related to Fig. 7.**

**a** Normalized stop codon ratio per testis (triangle) and ovary (circle) library. Codon frequencies were first normalized by sequence composition in piRNA clusters, and ratio was calculated as mean frequency of stop codons vs mean frequency of all other U*nn* sequences. Signal shown on a $\log_2$ scale. Libraries with robust (red) or near-robust (yellow) stop codon enrichment are indicated. **b** Difference in stop codon ratio per testis (triangle) and ovary (circle) library following normalization. Stop codon ratio was calculated as the mean frequency of stop codons divided by the mean frequency of all other U*nn* codons either with or without prior normalization for sequence composition in the piRNA clusters (ratios after normalization are shown in (a) and without in Fig. 6c). Signal shown on a $\log_2$ scale. Libraries with robust (red) or near-robust (yellow) stop codon enrichment are indicated. **c-d** Scatterplots showing correlation between 1U ratio and stop codon ratio either before (c) or after (d) normalization to cluster composition. Shaded area represents a 95% confidence interval (CI) of the linear regression fit determined through residual bootstrap with 1000 replicates.

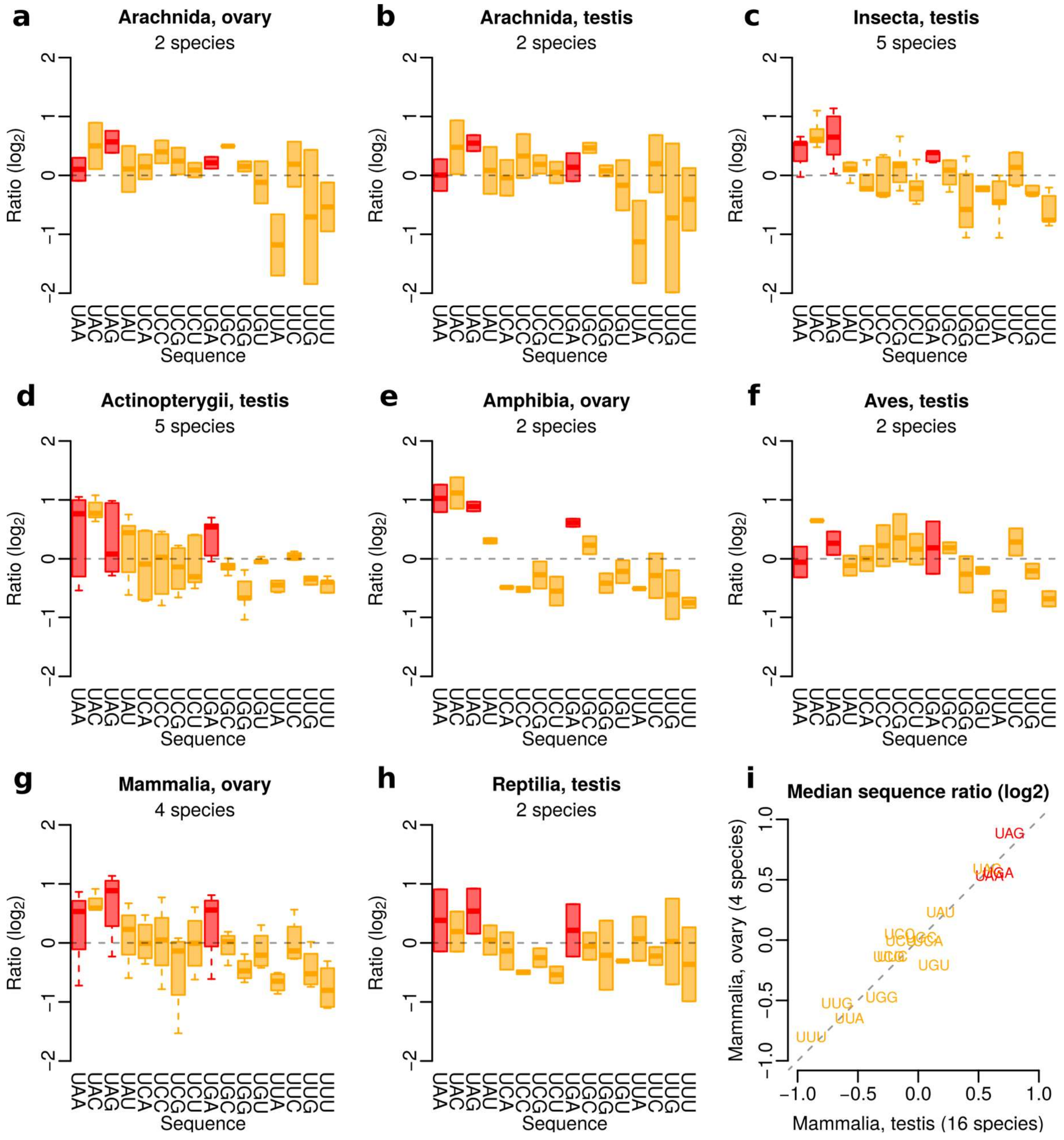


**Fig. S11. Related to Fig. 7.**

**a-h** Boxplots showing ratio between observed and expected fraction of U*nn* sequences in arachnid ovary (a), arachnid testis (b), insect testis (c), ray-finned fish testis (d), amphibian ovary (e), bird testis (f), mammalian ovary (g), and reptilian testis (h). Each species is represented as one data point in the boxplots, and multiple libraries from the same species were averaged. Boxplots show median (central line), interquartile range (IQR, box), and minimum and maximum values (whiskers, at most 1.5*IQR). **i** Scatterplot comparing the median sequence ratio across mammalian species with ovary (Fig. S11g) and testis (Fig. 7e) libraries.

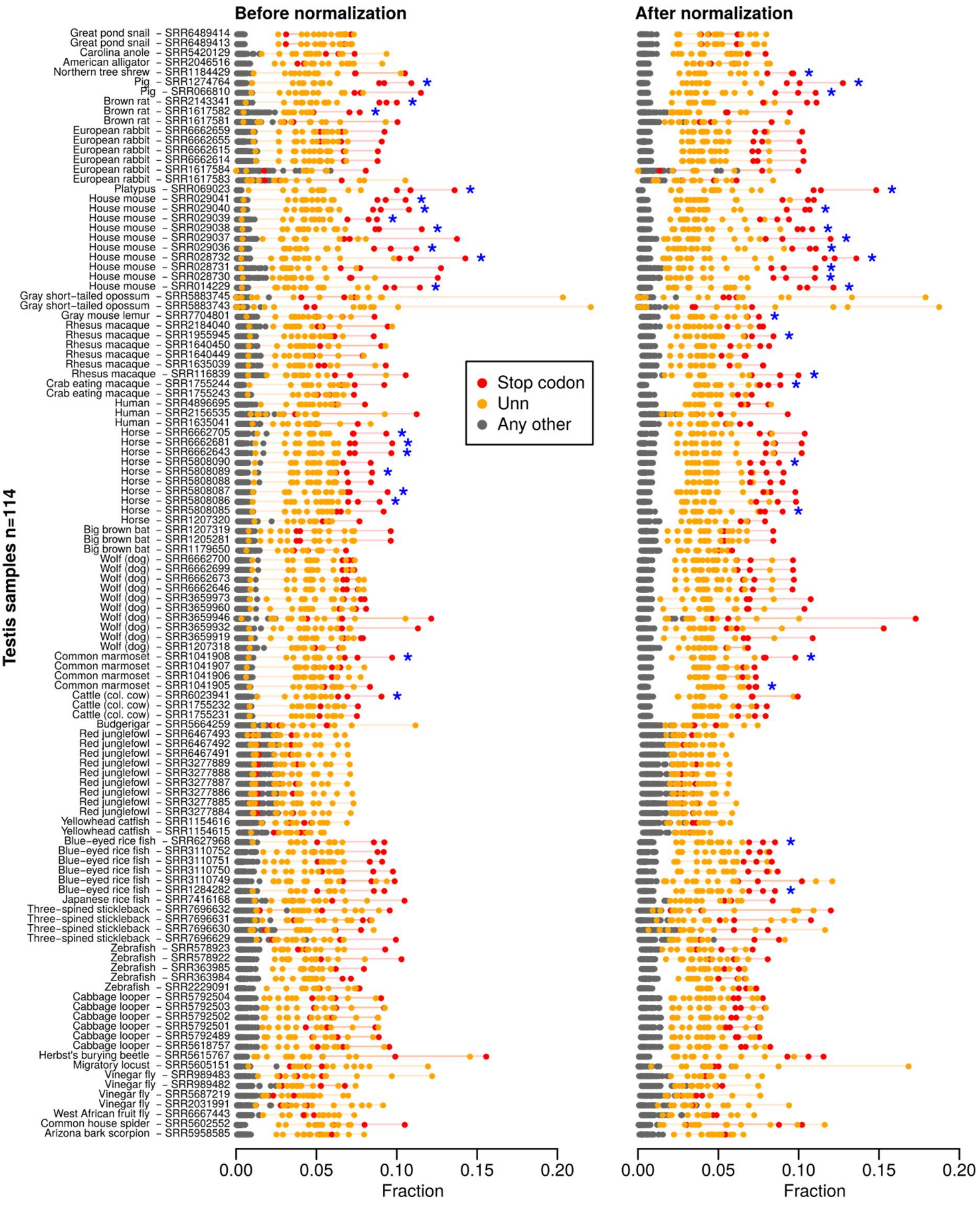


**Fig. S12. Overview of 114 libraries from testis. Related to Fig. 6 and Fig. 7.**
Detailed overview of 5' end sequences across all 114 libraries derived from testis before (left) and after (right) normalization to cluster composition. Each row represents one library, labelled by species and SRA accession code. All possible 5' end sequences are shown as circles and colour-coded to identify stop codons (*n*=3, red), other U*nn* sequences (*n*=13, orange), and all other sequences (*n*=48, grey). Horizontal lines connecting the red and orange circles, respectively, are displayed to enhance readability. Libraries with a robust stop codon enrichment are marked with a blue asterisk.

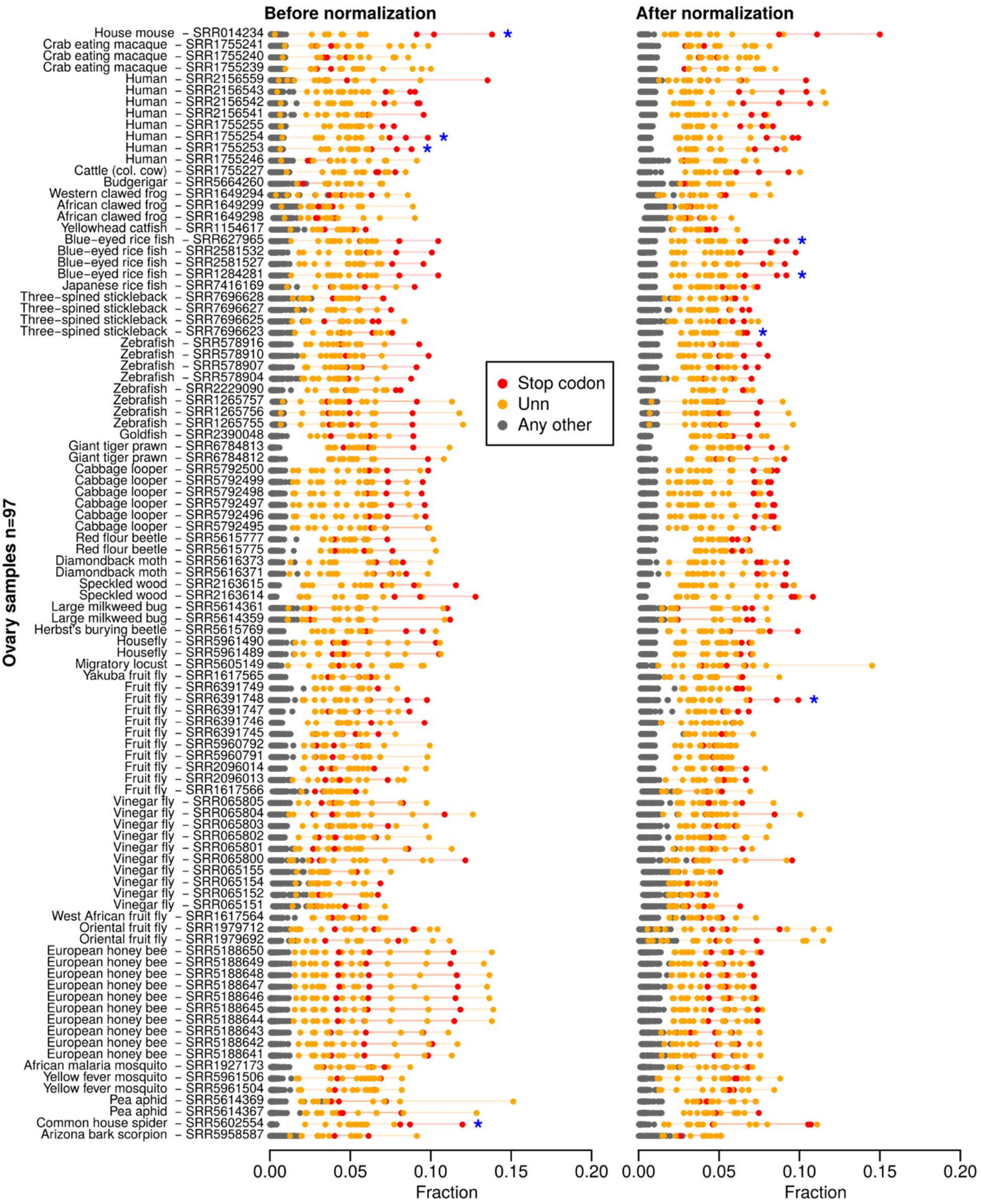


**Fig. S13. Overview of 97 libraries from ovary. Related to Fig. 6 and Fig. 7.**

Detailed overview of 5' end sequences across all 97 libraries derived from ovary before (left) and after (right) normalization to cluster composition. Each row represents one library, labelled by species and SRA accession code. All possible 5' end sequences are shown as circles and colour-coded to identify stop codons (*n*=3, red), other U*nn* sequences (*n*=13, orange), and all other sequences (*n*=48, grey). Horizontal lines connecting the red and orange circles, respectively, are displayed to enhance readability. Libraries with a robust stop codon enrichment are marked with a blue asterisk.

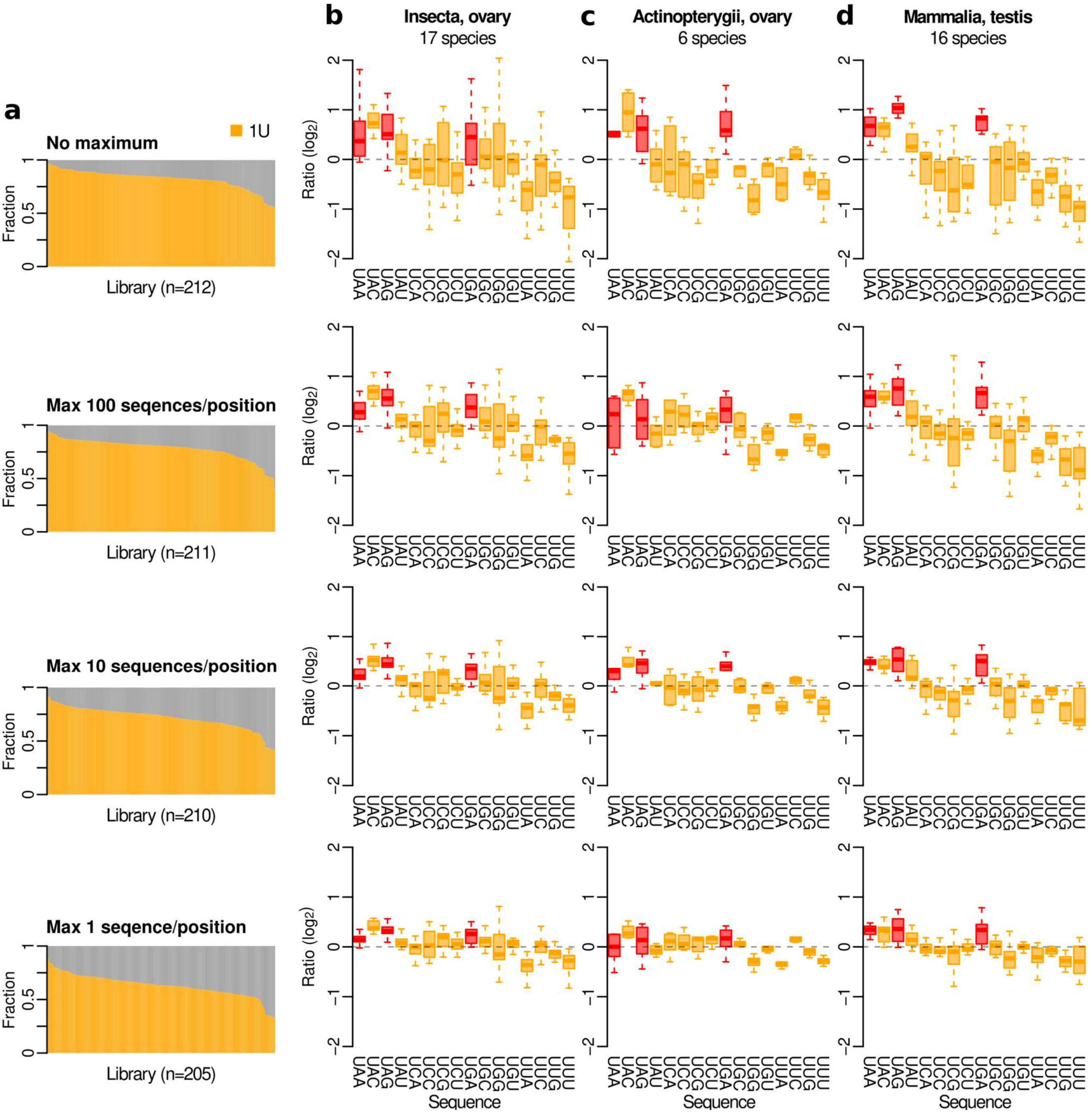


**Fig. S14. Stop codon enrichment can be detected using different subsampling thresholds.**
**a** Bar plots showing 1U signal across libraries from 49 species using four different levels of read subsampling as indicated by the headers. **b-d** Boxplots showing ratio between observed and expected fraction of U*nn* sequences in insect ovary (b), ray-finned fish ovary (c), or mammalian testis (d), using the same subsampling thresholds as in (a). Each species is represented as one data point in the boxplots, and multiple libraries from the same species were averaged. The three largest groups are shown. Please note that the "Max 100 sequences/position" filtering is the same one that was used throughout the manuscript (e.g., Fig. 6b, Fig. 7c-e).
